# varVAMP: automated pan-specific primer design for tiled full genome sequencing and qPCR of highly diverse viral pathogens

**DOI:** 10.1101/2024.05.08.593102

**Authors:** Jonas Fuchs, Johanna Kleine, Mathias Schemmerer, Julian Kreibich, Wolfgang Maier, Namuun Battur, Thomas Krannich, Somayyeh Sedaghatjoo, Lena Jaki, Anastasija Maks, Christina Boehm, Carina Wilhelm, Jessica Schulze, Christin Mache, Elischa Berger, Jessica Panajotov, Lisa Eidenschink, Björn Grüning, Markus Bauswein, Sindy Böttcher, Reimar Johne, Jürgen Wenzel, Martin Hölzer, Marcus Panning

## Abstract

Time- and cost-saving surveillance of viral pathogens is achieved by tiled sequencing in which a viral genome is amplified in overlapping PCR amplicons and qPCR. However, designing pan-specific primers for viral pathogens that have high genomic variability represents a major challenge. Here, we present a bioinformatics command-line tool, called varVAMP (<u>var</u>iable <u>v</u>irus <u>amp</u>licons). It relies on multiple sequence alignments of highly variable virus sequences and enables automatic pan-specific primer design for qPCR or tiled amplicon whole genome sequencing.

The varVAMP software guarantees pan-specificity by two means: it designs primers in regions with minimal variability and introduces degenerate nucleotides into primer sequences to compensate for common sequence variations. We demonstrate varVAMP’s utility by designing and evaluating novel pan-specific primer schemes suitable for sequencing the genomes of SARS-CoV-2, Hepatitis E virus, rat Hepatitis E virus, Hepatitis A virus, Borna-disease-virus-1, and Poliovirus. Moreover, we established highly sensitive and specific Poliovirus qPCR assays that could potentially simplify current Poliovirus surveillance. Importantly, wet-lab and bioinformatic techniques established for SARS-CoV-2 tiled amplicon sequencing were readily transferable to these new primer schemes and will allow sequencing laboratories to extend their established methodology to other human pathogens.

## INTRODUCTION

In recent years, next-generation full-genome sequencing of viruses has become an irreplaceable method to track the evolution of viral pathogens, study outbreaks in the human population and animal kingdom, and identify novel zoonotic threats^1–3^. While metagenomic analyses enable the broad analysis of viromes and potentially identify novel pathogens^4^, a high genome coverage is required to sufficiently analyze the genomic makeup of a specific viral population in order to e.g. reconstruct viral intra-host evolution^5,6^. This can be achieved by prior virus cultivation or increased sequencing depth, which have drawbacks. Virus cultivation is not always successful and can lead to cell-culture adaptations^7^. Moreover, increased sequencing depth is costly and might still result in poor genome coverage^8^. Targeted sequencing approaches via PCR-tiling or DNA hybridization allow highly specific sequencing on smaller machines without prior pathogen cultivation^9,10^. Particularly, PCR-tiling, in which the viral genome is amplified in overlapping fragments, has gained popularity due to its cost-effectiveness, low input requirement, and simple library preparation. The most prominent viral amplicon schemes were developed for SARS-CoV-2 in early 2020 and have allowed the sequencing of millions of viral genomes during the pandemic^11,12^. However, such amplicon schemes often need to be updated to reflect evolutionary changes or they have not been developed at all for many viral pathogens. Therefore, quantitative real-time PCR (qPCR) remains the diagnostic gold standard for analyzing patient samples for the presence of a viral pathogen^13^.

In an optimal setting, tiled-sequencing and qPCR primer designs for viral pathogens should be pan-specific. This can be challenging for viruses with a high genomic variability and common insertions and deletions (INDELs) sites. Thus, primers have to be designed in conserved regions with minimal genomic variation and should not span INDELs. As potential primer target regions might still display sequence variation, degenerate nucleotides can be introduced into primer sequences to further broaden their binding capacity. Optimal pan-specific primers should target highly conserved regions while keeping degeneracy minimal. This problem, coined maximum coverage degenerate primer design (MC-DGD), is a trade-off between specificity and sensitivity^14^. Primer-specific parameters complicate MC-DGD as not all potential regions are also potential primer binding sites^15^. Notably, qPCR designs have even more constraints due to additional hydrolysis probe-specific parameters and a low Gibbs free energy change (ΔG) of the target region^16^.

Various commercial and open-source primer design applications are available and often utilize primer3 at their core to calculate various primer parameters^17^. However, many of these tools were developed for a particular primer design problem and each of them only addresses some of the previously mentioned problems^18^. Primalscheme is the gold standard for designing tiled primer schemes for viral full genome sequencing^10^. However, primalscheme only handles genomic variations up to a sequence divergence of 5%, precluding its use for viral pathogens with significantly higher sequence divergence, such as Hepatitis E virus (HEV) or Hepatitis A virus (HAV)^19^. Moreover, primalscheme does not introduce degenerate nucleotides into primer sequences, limiting or even abolishing the binding affinity if mutations are located in the primer binding site^20^. Degenerate primer design has been elegantly addressed by software packages like easyPAC or DegePrime^21,22^, but they are not suited for the automatic design of tiled or qPCR schemes. For qPCR primer design, there are only a few open-source projects like QuantPrime^23^, but most software is not open access and available through commercial companies. However, none of these applications address pan-specific primer design, and not all calculate ΔG, resulting in time-intensive manual primer and amplicon evaluation.

Here, we present the command-line tool varVAMP (<u>var</u>iable <u>v</u>irus <u>ampl</u>icons) that enables fully automated pan-specific degenerate primer design for single amplicons, tiled amplicon schemes and qPCR and was tailored to viral genomics. We show varVAMP’s utility by designing and testing pan-specific tiled and qPCR primer sets for SARS-CoV-2, HEV (*Paslahepevirus balayani*), ratHEV (*Rocahepevirus ratti*), HAV (*Hepatovirus A*), Borna-disease-virus 1 (BoDV-1, *Orthobornavirus bornaense*), and Poliovirus (*Enterovirus C*, PV) 1-3, that represent different levels of sequence variability.

## MATERIAL AND METHODS

### Software

varVAMP is an easy-to-use cross-platform command line tool that is available via PyPI, DOCKER, BIOCONDA, and the Galaxy platform. It enables primer design for a variety of molecular techniques, including single amplicons, tiled full genome sequencing, and qPCR. In the following, we describe the different algorithms and steps that have been implemented for varVAMP. The primer design pipeline only requires an already computed multiple sequence alignment as input. Importantly, all parameters are highly customizable via direct arguments or a config file. varVAMP performs the following major steps (Fig. 1a): (1) Automatic parameter selection based on the input alignment, (2) alignment masking, (3) consensus sequence generation, (4) potential primer region search, (5) evaluation for primer or qPCR probe suitability of digested kmers, and (6) amplicon scheme creation. The last step differs depending on the three different modes available for varVAMP: Single, tiled, and qPCR. To evaluate potential off-targets, varVAMP can use a BLAST database to predict off-target effects and preferentially selects amplicons without off-targets in the final amplicon scheme.

**Figure 1.**
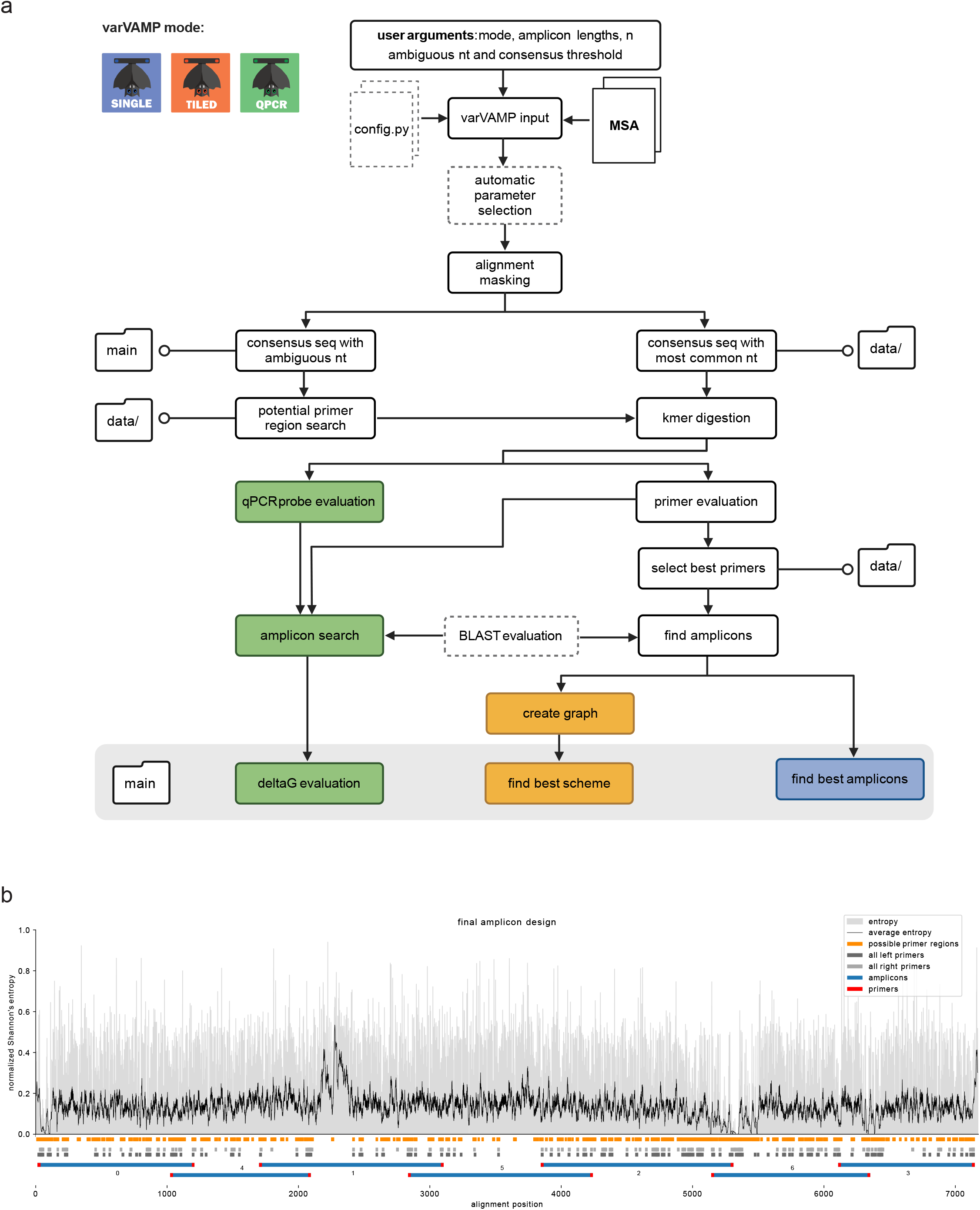
Schematic varVAMP overview and example output. **(a)** Overview of the varVAMP workflow. White boxes represent steps of the pipeline that are common to all modes. Consecutive steps are connected by arrows and optional steps are indicated with a dotted border. Colored boxes mark unique steps for each varVAMP mode (blue - single, orange - tiled, green - qPCR). Steps that produce outputs end in schematic folder icons for the main output and the additional data subfolder. (n - number, nt - nucleotide). **(b)** Example overview plot that is produced when running varVAMP. This plot was generated with varVAMP’s tiled mode on the example MSA of HEV-3 sequences provided as example data within the varVAMP github repository. Shown is the normalized Shannon’s entropy for each alignment position (gray) and its rolling average over 10 nucleotides (black curve). The orange boxes below the plot mark the start and stop MSA positions of potential primer regions (regions that have, in this case, a maximum of 4 ambiguous bases within the minimal primer length of 19) and the gray and light gray boxes mark all considered forward and reverse primers, respectively. The final scheme that was selected by the graph search for overlapping amplicons (blue) with low-penalty primers (red) is depicted at the bottom.

In a first step, varVAMP can estimate some of the user-parameters. For a minimal primer length *l*_*min*_, two main parameters influence the primer design: The number of ambiguous nucleotides tolerated within a primer sequence *n*_*a*_ with *n*_*a*_ *ϵ* ℕ, 0 ≤ *n*_*a*_ ≤ *l*_*min*_ and the identity threshold for a nucleotide *t* to be considered a consensus nucleotide with *t ϵℝ*, 0 ≤ *t* ≤ 1. Optimization is only performed for *n*_*a*_ or *t*, the other parameter has to be set manually. If *n*_*a*_ and *t* are both not given, *t* is optimized and *n* = 2. For each optimization iteration, *n*_*a*_ or *t* are incremented by − 1 or 0. 1, respectively. To perform parameter selection, the highest nucleotide frequency at each alignment position is determined. For each optimization step, the lengths of nucleotide stretches that consist of nucleotides reaching the current threshold {*l*_1_, *l*_2_ … *l*_*m*_} are calculated. The coverage *c* of the given alignment that can be considered for potential primer regions is then estimated by:

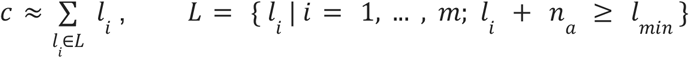

We define that optimization is reached if less than 50% of the alignment can be considered for potential primers. If varVAMP is used to design qPCR schemes, the number of ambiguous characters for the qPCR probe *n*_*p*_ is set to *n*_*p*_ = *n*_*a*_ – 1 to ensure a higher specificity of the probe compared to the flanking primers.

In the following preprocessing step, gaps in the alignment are masked. Given *n*_s_, the number of sequences in the alignment, common gaps are defined as gaps present in more sequences than *n*_s_ · (1 − *t*). Common gaps are then masked with ‘N’ (single nucleotide deletions) or ‘NN’ (larger deletions) in the multiple sequence alignment. This ensures that primers will not span regions that could be potential INDEL sites and that for the large majority of sequences, the amplicon size is not overestimated. Moreover, qPCR amplicons are not considered if they would span large deletions as small deviations in the amplicon length are particularly problematic for smaller qPCR amplicons.

In the next step, two consensus sequences are deduced from the gap-masked alignment. At each alignment position, the sorted list of observed nucleotide frequencies is calculated. If a sequence in the alignment contains ambiguous nucleotides, all permutations of these nucleotides are considered and added to the nucleotide frequencies proportionally to the number of permutations. The first consensus sequence is generated simply from the most frequent nucleotide at each site. This majority consensus sequence is the basis for the primer search. For the second consensus sequence, the observed nucleotide frequencies at each site are added up, starting from the highest frequency, until their sum reaches *t*. The IUPAC symbol for the set of nucleotides that contributed to the frequency sum is then taken as the consensus at the site. This symbol will be identical to the corresponding nucleotide in the majority consensus if the frequency of that nucleotide alone reaches or exceeds *t*. This second consensus sequence allows searching for regions that only have a certain amount of ambiguous characters within the minimal primer length.

Next, the consensus sequence with ambiguous nucleotide characters is searched for potential primer regions. The algorithm opens a region window at the start of the sequence. The window is closed if > *n*_*a*_ ambiguous nucleotides are found within a sequence of *l*_*min*_ nucleotides or a gap is reached. We define the resulting window as *w* = [*w*_*start*_, *w*_*end*_] with ambiguous character positions *x*_1_, …, *x*_*m*_. If the window was closed due to a gap, a new window is opened at the subsequent nucleotide *w*_*end*_ *+* 1. If the window was closed due to the number of ambiguous characters, the new window is opened at the position after the first ambiguous character counting towards *n*_*a*_ that led to closing the previous window, *x*_1_ *+* 1.Regions are only considered for the primer search if *w*_*end*_ − *w*_*start*_ ≥ *l*_*min*_.

In the identified primer search regions, the majority consensus is digested into all possible unique *k*-mers for *l*_*min*_ ≤ *k* ≤ *l*_*max*_. Afterwards, each kmer is evaluated for its suitability as a primer. For this primer3^17^ is used and some of the rationals and functions were adapted from primalscheme^10^. First, each kmer is hard-filtered independent of its direction for unacceptable temperature, size, GC content, homopolymer length, di-nucleotide repeats, and homodimer formation. Moreover and similar to primalscheme, a base penalty *p*_*b*_ for the kmers’ deviations from the optimal temperature, size and GC content is calculated and also hard-filtered if it exceeds the base penalty threshold. Then primers are evaluated for their suitability as forward or reverse primers by filtering out unacceptable hairpin formation temperatures, the 3-prime presence of ambiguous characters, and the absence of a GC clamp. For primers surviving all filtering steps, a permutation penalty *p*_*p*_ and a 3’ mismatch penalty *p*_*m*_ are calculated. *p*_*p*_ is calculated as the number of primer permutations of the primer version that has the ambiguous characters (deduced from the ambiguous consensus sequence) multiplied by the permutation penalty. For a primer with *n*_*p*_ characters, we define position-specific penalties 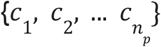 and calculate the mismatch frequencies at each position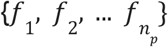. We then calculate:

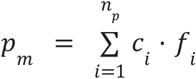

Note that only the last 5 positions receive non-zero multipliers in the standard settings. The final primer penalty *p*_*primer*_ is then calculated as:

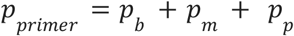

*p*_*primer*_ reflects the primers’ deviations from base parameters, its number of permutations and number of mismatches at the 3’-prime end. The closer *p*_*primer*_ is to zero, the better it represents an optimal primer.

To reduce the number of potential primers, all primers are penalty-sorted from low to high. From this sorted list, primers are retained if they do not overlap with the middle third of an already retained lower scoring primer, improving a final selection of primers with minimal overlap and minimal *p*_*primer*_. Next, all potential non-dimer forming combinations of forward and reverse primers within a given amplicon range length are computed. A resulting amplicon *a*_*i*_ is defined by primers *primer*_*fw*_, *primer*_r*v*_. Given the length of the amplicon *l*_*i*_ and the user-defined optimal amplicon length *l*_*opt*_, we define the amplicon penalty *p*_*i*_ :

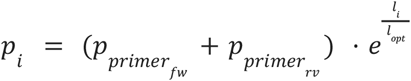

This ensures that the amplicon selection is length dependent and that it favors amplicons with a length closer to the optimal amplicon length. For a single-amplicon design, amplicons are sorted by their penalties from low to high and only low-scoring non-overlapping amplicons are retained.

For the tiled approach, a weighted directed graph *G* = (*V, E*) with vertices *V* and edges *E* is created. Each vertex *v*_*i*_ ∈ *V* represents an amplicon *a*_*i*_ and the set of vertices is given by *V* = {*v*_1_, *v*_2_, … *v*_*m*_}. We define the vertex start 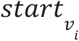 as the position of the first nucleotide in *primer*_*fw*_ belonging to *a*_*i*_ and the vertex stop 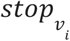 as the position of the last nucleotide in *primer*_*fw*_ belonging to *a*_*i*_. An edge e ∈ *E* is defined as a tupel *v*_*i*_, *v*_*j*_, *w*_*j*_ of two distinct nodes. The edge weight *w*_*j*_ = (*o*_*j*_, *p*_*j*_) incorporates the information about whether or not amplicon *a*_*j*_ generated an off-target hit with the optional BLAST database and amplicon penalty *p*_*j*_. If an off-target is generated *o*_*j*_ is 1 otherwise it is 0. The set of all edges *E* is defined as *E* = {(*v*_*i*_, *v*_*j*_, *w*_*j*_) | *v*_*j*_, *v*_*j*_ ∈ *V* and *i* ≠ *j* and *v*_*i*_, *v*_*j*_ overlap}. We say that *v*_*i*_, *v*_*j*_ overlap if they satisfy the user-defined reciprocal pairwise sequence overlap and 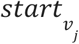 is not located in the first half of *v*. Next, varVAMP searches for the shortest path in *G* from a source vertex *v*_s_ with Dijkstra’s algorithm^24^. The stop position with the highest genomic index of all *v*_*i*_ ∈ *G* is denoted 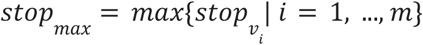 and the lowest penalized amplicon with the furthest stop position reached by Dijkstra’s search is termed *v*_*max*_. The amplicon coverage over the consensus sequence is defined as 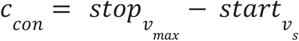. We store the current highest coverage 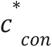 and the shortest path search is repeated for all *v*_*i*_ until 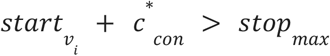. Therefore, the shortest path resulting in the highest coverage is the path that resulted in 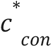. varVAMP defines two amplicon pools containing non-adjacent amplicons for the final scheme to allow primer multiplexing. In the last step, both pools are analyzed for the presence of primer heterodimers. If heterodimers are found, varVAMP considers the previously excluded primers overlapping with the middle third of the heterodimer-forming pair and tries to find primers that do not form heterodimers within the respective primer pools.

For the qPCR mode, the consensus sequence containing ambiguous nucleotides is searched for regions that satisfy the qPCR probe specific length and *n*_*p*_ constraints and is again digested into all possible unique kmers within the probe size range. These kmers are tested and evaluated for their suitability as qPCR probes in a manner analogous to primer screening. However, here we apply additional constraints: (i) probes are not allowed to have ambiguous bases at either end, (ii) probes cannot have a guanine at the 5’ end as this might result in quenching, and (iii) their direction is defined so that the qPCR probes have more cytosines than guanines. Next, varVAMP searches for potential qPCR amplicons. This is achieved by searching for primer subsets within the amplicon length constraint flanking a qPCR probe. Potential amplicons are excluded if they violate the GC content constraint or contain large deletions. The flanking primers must be within a narrow temperature range, the probe has to have a higher temperature than the primers, they cannot form dimers with each other, and the probe has to be within a certain distance to the primer on the same strand. varVAMP also evaluates the presence of dimers in all probe-primer permutations and excludes primer-probe combinations that overlap at their ends, as this might also lead to unspecific probe hydrolysis. Lastly, amplicons are sorted by their amplicon penalty 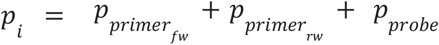 and tested for their ΔG at the lowest primer temperature using seqfold (https://github.com/Lattice-Automation/seqfold). varVAMP reports amplicons that pass the ΔG cutoff.

varVAMP has an optional BLAST feature that allows evaluation if amplicon primers could result in off-target products using a custom BLAST database^25^. Here, we perform a relaxed BLAST search with the BLAST settings published for primerBLAST^26^. Afterwards, the results are filtered for matches with a user-definable minimal overlap of identical nucleotides considering both query coverage and mismatches. We now check each amplicon for potential off-target hits defined as matches for both primers that are sufficiently close together on the same reference sequence, but on opposite strands. Amplicons that result in off-target hits, are preferentially not considered in the final scheme. In the single and qPCR mode, amplicons are first sorted for the absence of off-targets and then by their penalty. In the tiled mode, the shortest path is first evaluated on the amount of off-target hits generated by the path before considering the cumulative amplicon penalty, thereby avoiding, but not excluding amplicons with off-target effects.

The final primers (independently from the varVAMP mode) are deduced from the consensus sequence incorporating degenerate nucleotides.

### Primer design for the individual pathogens

Data selection for the multiple sequence alignments (MSAs) that were used as the inputs for varVAMP were highly dependent on the individual pathogens.

For SARS-CoV-2, we obtained 920,323 full-length genome sequences, sampled between 2021-10-11 and 2023-09-26, and their lineage assignments from https://github.com/robert-koch-institut/SARS-CoV-2-Sequenzdaten_aus_Deutschland (accessed 2023-10-13). Covsonar (https://github.com/rki-mf1/covsonar, v1.1.9) was used to calculate mutation profiles for all sequences, followed by a Python script (https://github.com/rki-mf1/sc2-mutation-frequency-calculator, v0.0.2-alpha) to select characteristic mutations per lineage (75% frequency). The script then uses these characteristic mutations to construct a single, representative consensus sequence per lineage. A representative consensus sequence was only calculated if at least ten genomes were available for a particular lineage. This resulted in representative consensus genomes for 865 SARS-CoV-2 lineages which were then used as input for varVAMP.

For BoDV-1 we downloaded all available full-length sequences that belong to the *Orthobornavirus bornaense* species (BoDV-1: 54 sequences, BoDV-2: 1 sequence). For HAV we downloaded all available full-length sequences that belong to the *Hepatovirus A* species (326 sequences). Patent and artificial clone sequences were excluded resulting in 309 HAV sequences. PV sequences were filtered in a similar manner and we excluded, by manual alignment inspection, highly divergent sequences that were likely the result of recombination events with other Enteroviruses yielding 944 sequences. For qPCR designs, we split this dataset, based on metadata, into the individual serotypes 1-3 resulting in 241, 494 and 209 sequences, respectively. For HEV data selection, we downloaded all available full-length sequences of the *Hepeviridae* family (1377 sequences). Patent and artificial clone sequences were excluded resulting in 1349 sequences. The remaining sequences were compared to the HEV reference set by Smith et al. 2020^27^, extended with the reference sequences for rat, bird, bat, fish, frog and planthopper HEV (NC_038504.1, NC_023425.1, NC_018382.1, NC_015521.1, NC_040835.1 and NC_040710.1, respectively), using the ggsearch36 algorithm^28^. Classification resulted in 1222 HEV sequences and 71 ratHEV sequences. Next, we used the greedy clustering algorithm of vsearch 2.22.1^29^ with global clustering thresholds of 0.82 and 0.71, respectively, to further split the HEV and ratHEV datasets by similarity. Clustering results were manually inspected in phylogenetic trees constructed with IQ-TREE 2 under the GTR+F+R10 substitution model and 1000 bootstrap replicates^30^. For HEV we choose two clusters that reflect the most common European HEV-3 subgenotypes (HEV-3 f, e and HEV-3 c, h1, m, i, uc, l). For ratHEV we focused on the largest cluster essentially excluding ratHEV from non-rat species and further excluded sequences that were too short resulting in a total of 41 sequences.

Next, the pairwise sequence identity within each sequence batch was calculated with Identity (https://github.com/BioinformaticsToolsmith/Identity)^31^ and the sequences were aligned with MAFFT^32^ with default settings. These alignments were then used as the input for varVAMP. Based on the sequence identity, we chose, for tiled sequencing, to fix the allowed max number of ambiguous characters within the minimum primer length depending on mean sequence identity within the batch (*n*_*a*_ = 2 with 90% identity, *n*_*a*_ = 4 between 70 and 80%, *n*_*a*_ = 5 below 70%). Next, the identity threshold *t* was maximized until varVAMP could not find a tiled scheme that covered the whole genome (Table 1). For the qPCR designs we chose to allow one less ambiguous base for the probe compared to the primers (Table 2). Here, settings were selected based on if varVAMP was able to find a qPCR scheme under the ΔG constraints rather than solely on sequence similarity. All input alignments and varVAMP outputs are available at: https://github.com/jonas-fuchs/ViralPrimerSchemes.

**Table 1.** Summary of varVAMP designs for tiled sequencing. The table lists important alignment statistics, varVAMP input parameters and output information including the varVAMP version. Pairwise sequence identity was calculated with Identity (https://github.com/BioinformaticsToolsmith/Identity). All primers and respective varVAMP outputs are accessible at: https://github.com/jonas-fuchs/ViralPrimerSchemes.

| alignment statistics |  |  |  | varVAMP parameters |  |  |  | varVAMP output |  |  |
| --- | --- | --- | --- | --- | --- | --- | --- | --- | --- | --- |
| virus | subtypes | n sequences | % mean sequence identiy | max ambig bases | threshold | optimal amplicon size | maximum amplicon size | expected recovery | n amplicons | varVAMP version |
| SARS-CoV-2 | B.1 - XBB | 865 | 99 ± 1 | 1 | 0.99875 | 700 | 800 | 99.72 % | 55 | v.0.9.4 |
| BoDV-1 | all | 55 | 89 ± 8 | 2 | 0.94 | 400 | 550 | 98.6 % | 27 | v.0.6 |
| HAV | all | 309 | 81 ± 10 | 4 | 0.93 | 1000 | 1600 | 95.65 % | 7 | v.0.8.3 |
| HEV genotype 3 | f, e | 376 | 76 ± 6 | 4 | 0.91 | 1000 | 1500 | 99.02 % | 7 | v.0.8.2 |
| HEV genotype 3 | c, h1, m, i, uc, l | 201 | 75 ± 9 | 4 | 0.90 | 1000 | 1500 | 99.28 % | 6 | v.0.8.2 |
| PV | 1-3 | 944 | 71 ± 13 | 4 | 0.91 | 1000 | 1400 | 99.63 % | 7 | v.0.8 |
| ratHEV | all | 41 | 57 ± 10 | 5 | 0.82 | 1200 | 1700 | 97.41 % | 6 | v.0.8.3 |

**Table 2.** Summary of varVAMP designs for qPCR. Important alignment statistics are listed together with varVAMP input parameters and output information including the varVAMP version. Pairwise sequence identity was calculated with Identity (https://github.com/BioinformaticsToolsmith/Identity). All primers and respective varVAMP outputs are accessible at: https://github.com/jonas-fuchs/ViralPrimerSchemes.

| alignment statistics |  |  |  | varVAMP parameters |  |  |  | varVAMP output |  |
| --- | --- | --- | --- | --- | --- | --- | --- | --- | --- |
| virus | subtypes | n sequences | % mean sequence identiy | primer max ambig bases | probe max ambig bases | threshold | $\Delta G$ cutoff (kcal/mol) | n found schemes | varVAMP version |
| PV | 1 | 241 | 86 ± 12 | 2 | 1 | 0.93 | -3 | 3 | 0.7 |
| PV | 2 | 494 | 88 ± 8 | 1 | 0 | 0.98 | -3 | 4 | 0.7 |
| PV | 3 | 209 | 91 ± 9 | 2 | 1 | 0.93 | -3 | 3 | 0.7 |

### HEV qPCR and HEV sub-genotyping

Patient serum samples were tested for HEV using the HEV RT-PCR Kit 1.5 (AS0271543, Altona Diagnostics, AltoStar®). For HEV sub-genotyping of HEV-positive samples, we used an in-house nested RT-PCR (210212, Qiagen, Hilden, Germany) protocol based on previously published primers and nested primers in a conserved region of ORF1^33^. RT-PCR was performed at 42 °C for 60 min, 15 min at 95°C followed by 40 cycles at 94 °C (30 s), 56.5 °C (30 s) and 74 °C (45 s). Final elongation was at 74 °C for 5 min. Agarose gel negative PCR reactions were subjected to a nested PCR reaction. Afterwards, PCR products were Sanger sequenced.

### Production of virus stocks

The BoDV-1 strains were derived from native human brain sections^34–36^ and were propagated in Vero cells, which were grown in DMEM supplemented with 10% heat-inactivated fetal calf serum (FCS), 90 U/ml streptomycin, 0.3 mg/ml glutamine, and 200 U/ml penicillin (all PAN Biotech, Aidenbach, Germany). Permanently infected cells were split twice a week and monitored for *Mycoplasma* spp. contamination every 12 weeks. To obtain a cell-free viral stock, cell culture supernatants were centrifuged at 1,000 rcf to remove cell debris and filtered through Rotilab syringe filters with a pore size of 0.22 μm (Carl Roth, Karlsruhe, Germany).

HEV-containing supernatant was harvested from persistently infected cell culture. The liver carcinoma cell lines (PLC/PRF/5, ATCC: CRL-8024) persistently infected with HEV-3c strain 14-16753, HEV-3e strain 14-22707 or HEV-3f strain 15-22016 (provided by National Consultant Laboratory for HAV and HEV, University Hospital Regensburg) were maintained in modified Minimum Essential Medium at 37 °C and 5% CO_2_^37^.

HAV genotype IB strains MBB^38^ and V18-35519 (derived from plasma of a patient with acute hepatitis A) were both propagated in HuH-7 cells maintained in BMEM and incubated at 34.5 °C and 5% CO_2_. MBB and V18-35519 strains were harvested at 665 and 378 days post inoculation, respectively.

RD-A cells were infected with PV for virus propagation and cultivated in MEM Earls media with L-glutamine, 1x Non-essential amino acids, 100 U/ml penicillin and streptomycin (61100087, 11140050, 15140122, Thermo Fisher Scientific, Germany) and 7.3% heat inactivated fetal calf serum (BioWest, South America). RD-A cells were split once a week and an internal quality control performed at passage five to ensure cell sensitivity. The cell culture was conducted at 37 °C and 5% CO_2_. Cultures were checked daily for a cytopathic effect (CPE) for a maximum of seven days. Up to 2 ml of cell cultures with an observed CPE were centrifuged 4 min at 4000 rpm and the supernatant used as viral stock.

RatHEV strain R63^39^, which was originally detected in a Norway rat from Germany, and ratHEV strain pt2^40^, which was identified in a human patient in Hong Kong, were generated and propagated in HuH-7-Lunet BLR cells under conditions as described previously^41,42^. The ratHEV positive culture supernatants were harvested after 66 days post infection

### Tiled Illumina sequencing for HEV-3

Viral RNA was isolated using the QIAamp® Viral RNA kit (52904, Qiagen, Hilden, Germany) following the manufacturer’s protocol. Subsequently, a one-step RT-PCR using the SuperScript™ IV One-Step RT-PCR System (12594025, Thermo Fisher) was performed for each amplicon separately with a total primer concentration of 1 μM. To reduce non-specific amplification, reverse transcription was performed at 55 °C for 60 min and we gradually reduced the primer annealing temperature during the PCR in the first 10 cycles (10 sec at 98 °C, 10 sec at 63 °C (−0.5°C/cycle), 2 min at 72 °C) and then performed another 35 cycles at a constant annealing temperature (10 sec at 98 °C, 10 sec at 58 °C, 2 min at 72 °C). Next, amplicons were pooled and purified using AMPure XP beads (A63881, Beckman Coulter). 50-100 ng of DNA were prepared for Illumina sequencing using the NEBNext Ultra II FS DNA Library Prep Kit (E6177, NEB, Frankfurt am Main, Germany). Normalized and pooled sequencing libraries were denatured with 0.2 N NaOH and sequenced on an Illumina MiSeq instrument using the 300-cycle MiSeq Reagent Kit v2 (MS-102-2002, Illumina).

### Tiled Illumina sequencing for BoDV-1, HAV, ratHEV

Viral RNA was isolated using the QIAamp Viral RNA Mini Kit (Qiagen, Hilden, Germany) or the EMAG Nucleic Acid Extraction System (Biomeriéux Deutschland GmbH, Nürtingen, Germany). RNA was transcribed into cDNA with LunaScript RT SuperMix Kit (New England Biolabs, Ipswich, MA, USA) for 2 min at 25 °C, 10 min at 55 °C and 1 min at 95 °C. The cDNA was then amplified in single reactions with primer pairs, as well as multiplex PCRs with primer pools using the Q5 Hot Start-Fidelity DNA Polymerase Kit (New England Biolabs, Ipswich, MA, USA) with an initial step at 98 °C for 30 sec, followed by 35 cycles (15 sec at 98 °C, 5 min 65 °C) and a final extension step at 65 °C for 5 min. Illumina sequencing was performed analogous to the HEV-3 sequencing protocol.

### Tiled Illumina sequencing for PV

PV vaccine strain 1-3 (Sabin 1-3) RNA was isolated using the QIAamp® Viral RNA kit (52904, Qiagen, Hilden, Germany) following the manufacturers’ protocol from infected RD-A cells, with PV2 being archived RNA due to containment reasons. Subsequently, a one-step RT-PCR using the Qiagen One-step RT-PCR kit (210212, Qiagen, Hilden, Germany) was performed for each amplicon separately or for the respective pools with a total primer concentration of 0.6 μM. Reverse transcription step was done for 30 min at 50 °C followed by an initial PCR activation step at 95 °C for 15 min. Product amplification was done in 40 cycles with a stepwise reduction of the primer annealing temperature during the first 10 cycles (30 sec at 94 °C, 45 sec at 70 °C (ΔT −1 °C/cycle), 90 sec 72 °C) and a constant annealing temperature for the next 30 cycles (30 sec at 94 °C, 45 sec at 60 °C, 90 sec 72°C) and a final extension step for 10 min at 72 °C. Multiplex RT-PCR pools of each sample were combined and purified using MagSi-NGSPREP-PLUS beads (MDKT00010075, Steinbrenner, Germany) according to the manufacturer’s manual and DNA concentration measured using the Qubit™ 1X dsDNA Assay-Kit (Q33230, Thermo Fisher Scientific, Germany).

For Illumina sequencing, library preparation was done using 70-400 ng DNA with the Nextera XT DNA Library Preparation Kit (FC-131-1096, Illumina) and sequenced on an Illumina MiSeq Instrument (2 x 300 bp read length).

### Tiled ONT sequencing for HAV

Nucleic acid was extracted from samples on an EZ1® Advanced XL workstation using the EZ1 Virus Mini Kit v2.0 (Qiagen, Hilden, Germany) and transcribed into cDNA with LunaScript RT SuperMix Kit (New England Biolabs, Ipswich, MA, USA) for 2 min at 25 °C, 10 min at 55 °C and 1 min at 95 °C. The cDNA was then amplified in multiplex PCRs with HAV-specific primer pools using the Q5 Hot Start-Fidelity DNA Polymerase Kit (New England Biolabs, Ipswich, MA, USA) with an initial step at 98 °C for 30 sec, followed by 35 cycles (15 sec at 98 °C, 5 min 65 °C) and a final extension step at 65 °C for 5 min. Barcoding was performed with eight samples per run using the Rapid Barcoding Kit 96 V14 (Oxford Nanopore Technologies, Oxford, UK). The library was sequenced on an Mk1C (MinKNOW software version 23.07.12) for 72 hours using an R10.4.1 Flow Cell.

### Tiled ONT sequencing for SARS-CoV-2

A total of 14 SARS-CoV-2 positive extracts, tested at the Robert Koch Institute were selected. These samples originated from IMSSC2-lab networks that were received under the RKI integrated genomic surveillance program. All samples were nasopharyngeal or oropharyngeal swabs originating from patients in Germany during January-February 2024. Total nucleic acid extraction was done using MagNA Pure 96 DNA and viral NA Small Volume kit (Roche Life Science, Mannheim, Germany) on an automated extraction instrument (MagNA Pure 96 system, Roche Diagnostics) according to the manufacturer’s manual. Reverse-transcription was performed on viral RNA extracts using LunaScript® RT SuperMix (New England Biolabs, as part of the NEBNext ARTIC SARS-CoV-2 Companion Kit (Oxford Nanopore Technologies) according to the manufacturer’s protocol. Amplification was performed with 35 cycles of annealing temperature of 60 °C for 2 min and elongation at 72 °C for 3 min for both pools, without amplicons cleanup afterward. The barcoding was done using ONT Native Barcoding Expansion kit (EXP-NBD196). Fourteen samples together with 2 negative controls were multiplexed on a FLO-MIN 114 flow cell version R10 and sequenced on a GridION Mk1 device for 16 hours.

### RT-qPCR (PV 1-3)

The different varVAMP RT-qPCR assays were performed on serial dilution series of PV 1-3 (Sabin) RNA with a RNA concentration ranging from 1-3 ng/μl of the stock solution to evaluate the performance and sensitivity of the different assays. For proof of lack of cross-detection between the PV vaccine strains, all possible combinations were tested. Quantitative realtime RT-PCR was carried out using the 4X CAPITAL™ 1-Step qRT-PCR Probe Master Mix (BR0502002, Biotechrabbit, Germany) on a Roche instrument, following the manufacturer’s instructions. The reaction contained 0.4 μM of each primer with a total reaction volume of 20 μl. qRT-PCR cycling program started with reverse transcription at 50 °C for 10 min followed by an activation step at 95 °C for 3 min and 45 cycles for target amplification (95 °C for 10 sec, 59 °C for 30 sec). Primer annealing temperature was chosen after preliminary tests with different annealing temperature settings ranging from 56 - 64 °C on a Biometra TAdvanced (analytik jena, Germany) and product observation on an 1.5% agarose gel. The fluorescence was measured during the extension step with three different channel setups respective to the probe fluorophore (FAM = 465-510 nm, JOE = 533-580 nm, CY5 = 618-660 nm).

### Sequencing data analysis

The de-multiplexed raw Illumina reads were subjected to a custom Galaxy pipeline which we had initially developed for tiled amplicon sequencing of SARS-CoV-2^6^. These reads were pre-processed with fastp (v0.20.1) ^43^ and mapped to respective closely related genomes using BWA-MEM^44^ (v0.7.17). Importantly, the 3’ and 5’ regions of the viral reference genome were masked prior mapping until the 5’ and 3’ end of the flanking primer binding regions, respectively, as no novel information can be generated in these regions. Post mapping, primer sequences were trimmed with ivar trim (v1.3.1). Variants (SNPs and INDELs) were called with the ultrasensitive variant caller LoFreq^45^ (v2.1.5), demanding a minimum base quality of 30 and a coverage of at least 20-fold. Afterwards, the called variants were filtered based on a minimum variant frequency of 10% and on strand bias support. Finally, consensus sequences were constructed with bcftools (v1.15.1)^46^. Regions at both genome ends that lie outside the amplicons, regions with low coverage (<20x) or variant frequencies between 0.3 and 0.7 were masked with Ns.

For Oxford nanopore sequencing of SARS-CoV-2, poreCov (v1.9.3), a Nextflow workflow specifically tailored for SARS-CoV-2 genome reconstruction from nanopore amplicon data, was used to perform mapping (minimap2; v2.17)^47^, primer clipping, variant calling (Medaka; v1.8.0), and consensus genome reconstruction^48^. We ran poreCov to initially filter reads below 400 bp (--minLength 400) and above 1 kbp (--maxLength 1000) while coverage downsampling was disabled (--artic_normalize 0). The r1041_e82_400bps_sup_v4.2.0 model was used for variant calling with Medaka and detected variant calls filtered by a minimum base quality of 20 and a coverage of at least 20-fold. We set the allelic frequency of called mutations to 1 to ensure compatibility with the Illumina data for the in silico analysis.

For Oxford nanopore sequencing of HAV samples, pod5 raw data was duplex basecalled with Dorado version 0.5.3 and demultiplexed. Subsequent, fastq files were again processed using a custom Galaxy pipeline. First, raw data was pre-processed with fastp^43^ excluding reads <50 bp and >2000 bp. Afterwards, reads were mapped to the HAV reference genome NC_001607 using minimap2 (v2.17)^47^ and trimmed with ivar trim (v1.3.1). Variants were called with medaka (v1.3.2) and consensus sequences were constructed with bcftools (v.1.15.1)^46^. Regions at both genome ends that lie outside the amplicons and regions with low coverage (<20x) were masked with Ns.

### Data analysis and visualization

qPCR amplification curves were analyzed using the Roche LightCycler 480 II device software version LCS480 1.5.1.62 (Roche Applied Science). Data was analyzed using the 2nd derivative. Mapped bam files were analyzed and visualized with BAMdash v.0.2.4 (https://github.com/jonas-fuchs/BAMdash). We used GraphPad Prism 8 (genome recovery and qPCR), R 4.3.2 (phylogenetic tree) or python 3.11 (remaining figures) for data analysis and visualization. The data and code to reproduce the figures is available at: https://github.com/jonas-fuchs/varVAMP_in_silico_analysis. The schematic varVAMP workflow and data preparation workflow were created with biorender (https://www.biorender.com).

## RESULTS

### Software and output

The command-line tool varVAMP was written in python3 and requires only a pre-computed MSA as input. Notably, varVAMP is cross-platform (Windows 11, MacOS and Linux) with python 3.9 or higher being the single requirement prior to installation. varVAMP can design primers for single amplicons, tiled amplicon schemes and qPCR. The pipeline consists of multiple steps that are common to all different modes (alignment preprocessing, consensus generation and primer evaluation), mode-specific or optional (automatic parameter search and BLAST evaluation) (Fig. 1a). At its core, varVAMP wraps Primer3^17^ and uses a kmer-based approach to find all potential primers in a consensus sequence calculated from the input MSA. varVAMP addresses the MC-DGD problem by first calculating two consensus sequences that consist either of the majority nucleotides at each position or integrate degenerate nucleotides. The latter is used to find potential primer regions that are regions with a user-defined maximum amount of degenerate nucleotides within the minimal primer length. Afterwards, kmers of the majority consensus sequence that lie within these potential primer regions are tested for all relevant primer parameters. varVAMP evaluates these primers via a penalty system that incorporates information about primer parameters, 3’ mismatches, and degeneracy. In its tiled sequencing mode, varVAMP finds overlapping amplicons spanning the alignment while minimizing primer penalties by using Dijkstra’s algorithm for finding the shortest paths between nodes in a weighted graph^24^. For qPCRs, varVAMP evaluates probe and primer parameters independently and tests the ΔG of potential qPCR amplicons. The final primers are then deduced from the consensus sequence incorporating degenerate nucleotides. For some of the more computationally intensive tasks, the program is capable of using multicore processing. Although we have not extensively evaluated the running times, varVAMP typically finishes within seconds to minutes. This is highly dependent on the alignment’s size and number of sequences, as well as the alignment’s sequence variability that directly influences the amount of found primers. varVAMP produces multiple outputs in standardized formats and a plot displaying the alignment’s normalized Shannon’s entropy, all potential target regions, all primers that passed the initial filtering steps and the final amplicon design with low penalty primers (Fig. 1b).

### Design and evaluation of HEV pan-specific primers

HEV of the genus *Paslahepevirus* is the most common cause of acute viral hepatitis worldwide and is phylogenetically separated into four distinct genotypes (genotypes 1-4). In risk groups such as immunocompromised patients, the zoonotic HEV genotype 3 (HEV-3) can cause acute or chronic hepatitis^49^. HEV-3 has a high prevalence in industrialized countries and is further classified into subgenotypes with varying prevalence depending on the geographic region^50^. Most genome sequences show exceptional variability^51^ and have to be generated from the initial patient material as virus isolations require optimized cell culture systems^37^. To provide a simple sequencing procedure from patient material and to test varVAMP’s real-world applicability, we set out to design primers for HEV-3 tiled sequencing as a proof-of-principle. We initially downloaded all available full-genome HEV sequences from NCBI’s genbank and classified the (sub-) genotypes using fasta36 as previously described (Fig. 2a)^2^. Our aim was to design primers that would be specific for multiple HEV-3 sub-genotypes. Therefore, sequences were clustered based on their similarity using vsearch^29^ and the clustering result evaluated by constructing a maximum-likelihood phylogenetic tree with IQ-TREE 2^30^. Clustering resulted in seven clusters with more than 6 sequences (Fig. 2b). Four large clusters belonged to HEV-3 each comprising multiple subgenotypes. We decided to design primers for cluster 2 (HEV-3 f, e) and cluster 4 (HEV-3 c, h1, m, i, uc, l) to reflect the most common European HEV-3 subgenotypes^2^. Therefore, sequences of cluster 2 and 4 were separately aligned with MAFFT^32^ and the alignments used as the input for varVAMP yielding seven and six 1-1.5 kb amplicons, respectively (Fig. 2a and Table 1). Next, we evaluated these primer schemes on persistently HEV-3 f (strain: 15-22016) and c (strain: 14-16753) infected cell cultures using a one-step RT-PCR protocol. Agarose gel electrophoresis showed consistent and strong amplification for all amplicons with only a few unspecific bands for amplicon 2 and 3 of cluster 4 (Fig. 2c). Next generation Illumina sequencing of the pooled PCR products resulted in an even and high coverage for both samples (Fig. 2d). To further evaluate the primer schemes, we applied our protocol to a third HEV-3 e (strain: 14-22707) persistently infected cell culture for cluster 2 and to HEV-3 positive patient material for both clusters. In order to select the proper amplicon scheme, we first subclassified HEV-3 positive blood samples. Next, we evaluated the cluster 2 and 4 primer schemes on HEV-3 e (n=2) or HEV-3 c (n=4) samples, respectively. Next-generation sequencing results were comparable to the prior results and allowed HEV-3 genome reconstruction (Fig. 2e and S1). However, for patient 2 of cluster 2 and patient 4 of cluster 4 we observed a single amplicon dropout (S1). Interestingly, both dropouts were caused by amplicons with a forward primer close to the HEV-3 hypervariable region^51^ suggesting the presence of potential INDELs or variations that might have restricted primer binding.

**Figure 2.**
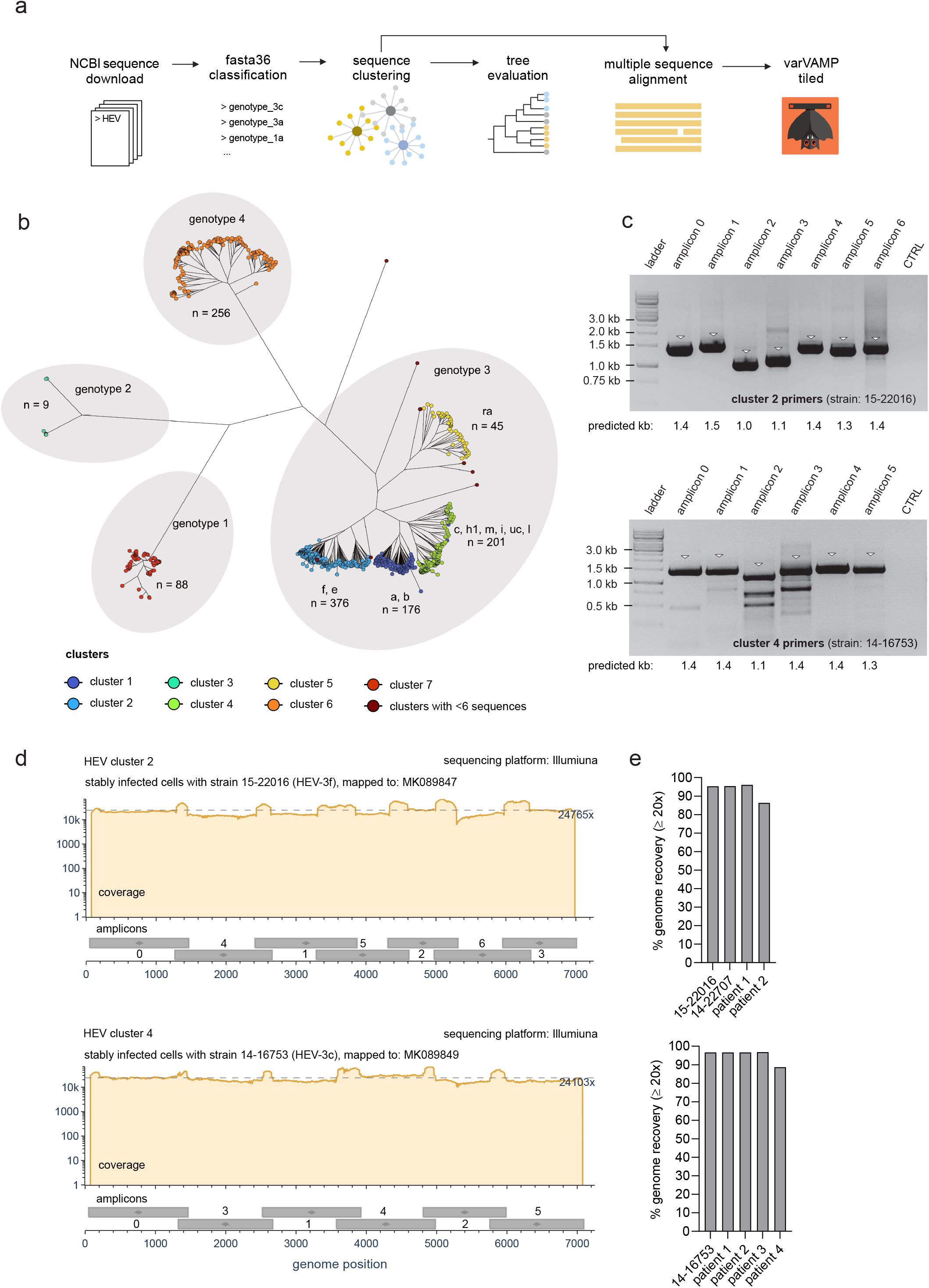
Primer design and tiled sequencing of HEV-3. **(a)** Schematic overview of the data preparation steps preceding primer design. All full-length sequences of HEV were downloaded from NCBI, sub-genotyped with fasta36 and clustered by similarity with vsearch. The clustering result was evaluated by phylogenetic tree construction. Afterwards, clusters comprising multiple subgenotypes were aligned with MAFFT and the MSA used as the input for varVAMP **(b)** Phylogenetic tree of full-length HEV sequences constructed with IQ-TREE 2 (GTR+F+R10, 1000 bootstrap replicates). The vsearch clustering results for each sequence is displayed in colors and the HEV genotypes and subgenotypes are indicated at the respective branches (n = number of sequences). **(c)** Agarose electrophoresis pictures of the individual PCR products for the cluster 2 (upper plot) and cluster 4 (lower plot) primer schemes tested with supernatant of HEV-3 f or HEV-3 c stably infected PLC/PRF/5 cells, respectively. Triangles indicate bands at the expected molecular weight of the PCR products (kb - kilobases) **(d)** Coverage plots of the Illumina sequencing results of the in (c) amplified PCR products for cluster 2 (upper plot) and cluster 4 (lower plot) mapped to their respective NCBI reference sequences MK089847 and MK089849. Below each coverage plot the genomic start and stop positions of each amplicon are displayed as gray boxes with their respective amplicon number. Dotted lines indicate mean coverages. Coverage plots were created with BAMdash (individual coverage plots are given in S1). **(e)** Genome recovery of HEV-3 persistently infected cell cultures and sub-genotyped HEV-3 positive blood samples that were subjected to their respective tiled amplicon workflow for cluster 2 (upper plot) or cluster 4 (lower plot). Genome recovery was calculated as % of reference nucleotides covered at least 20 fold. All PCR reactions were performed in the singleplex setting.

In summary, we used varVAMP to design two tiled primer schemes each specific for multiple HEV-3 sub-genotypes and used Illumina sequencing to recover near-to-complete viral genomes for both infected cell cultures and patient material.

### *In silico* design and evaluation of primer schemes for multiple viral pathogens with diverse sequencing variability

We designed primer schemes for tiled full-genome sequencing for SARS-CoV-2, BoDV-1, HAV, PV and ratHEV that display different degrees of sequence conservation over the whole genome with SARS-CoV-2 having the lowest (99 % pairwise identity) and ratHEV the highest overall sequence variability (57 % pairwise identity) (Table 1 and Fig. 3a). Similar to HEV-3, pan-specific amplicon sequencing protocols would massively simplify diagnostics and surveillance. The initial data selection and pre-processing was highly dependent on the individual data sets and inspired by our experiences with HEV-3. Only for SARS-CoV-2, we did not directly align sequences from public databases, but created representative consensus sequences of circulating lineages in Germany between October 2021 and September 2023 (920k samples) to represent the most prevalent variations for each lineage within the alignment. Depending on the mean pairwise identity, we chose to tolerate one to five ambiguous bases within the primer sequences and optimized the identity threshold (Table 1). With the exception of BoDV-1 and SARS-CoV-2, we aimed for an amplicon size of over 1000 bp so amplicons could span regions with an overall higher variability in which potential primers are scarce (Fig. 3a). Next, we evaluated the designed primer schemes *in silico* prior to wet-lab evaluation.

**Figure 3.**
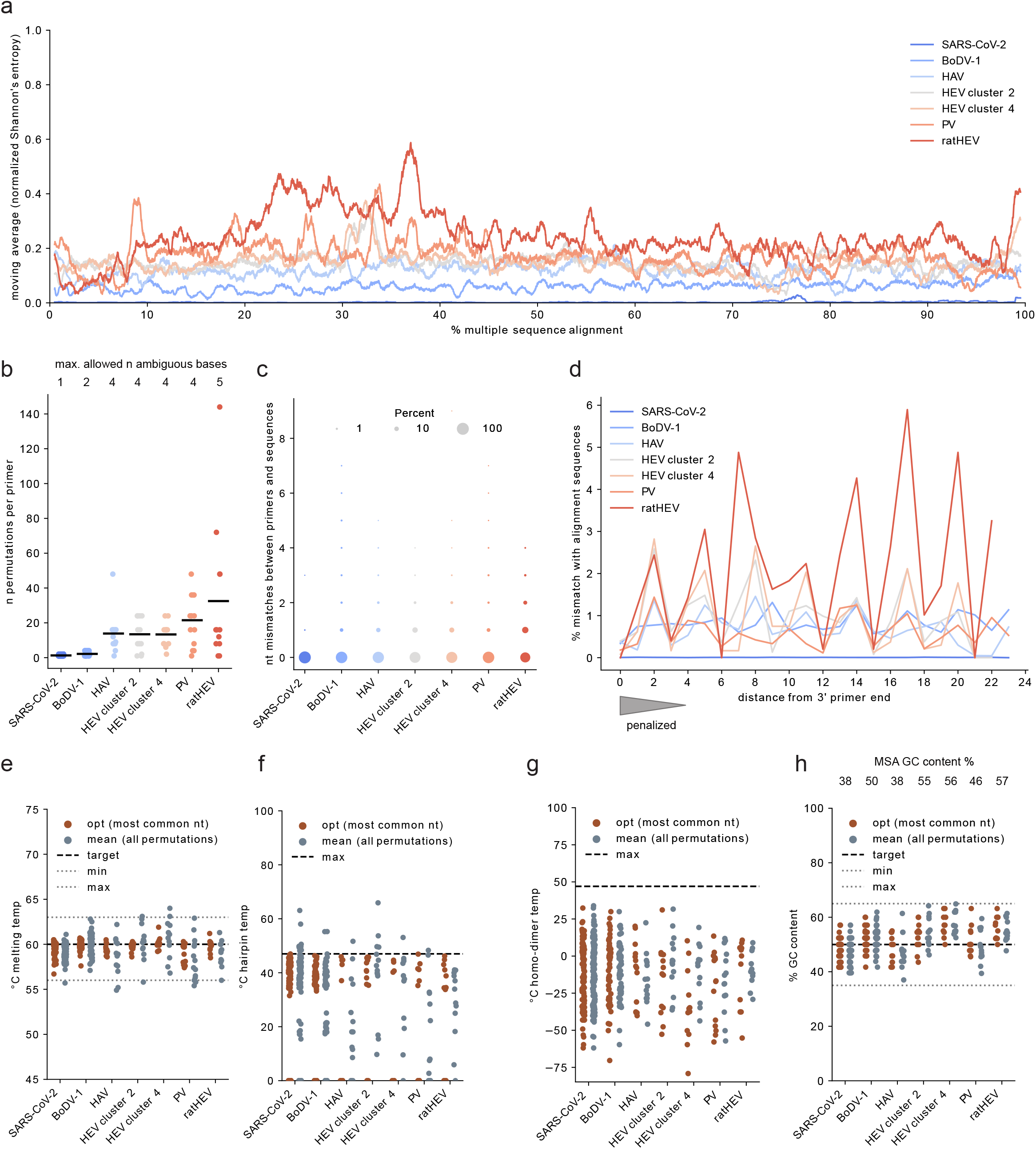
*In silico* evaluation of novel tiled primer schemes for SARS-CoV-2, BoDV-1, HAV, HEV, PV and ratHEV. **(a)** Normalized Shannon’s entropy (1% rolling average) of the MSA used as the varVAMP input. **(b)** Number of permutations (degeneracy) of each primer in the tiled sequencing scheme for the respective viruses. Each dot shows the degeneracy of a single primer. Horizontal lines indicate the mean. (n - number) **(c)** Cumulative counts of mismatches between primers and sequences in the varVAMP input MSA. For each primer the number of mismatches with each sequence of the MSA was counted if it was not covered by any primer permutation. Shown are the cumulative mismatches between primers and MSA sequences in the tiled primer schemes for the respective viruses. Dot area size is proportionate to the percentage. **(d)** Analogous to (c) the mismatches with the MSA sequences were counted per primer nucleotide position and averaged over all primers in a scheme. As primers vary in their length, the % mismatches are displayed starting at the primer’s 3’ end (position 0 is the most 3’ nucleotide position). The gray triangle schematically indicates the primer positions that varVAMP penalizes and the position-specific penalty multipliers (32, 16, 8, 4, 2). **(e-h)** Primer melting temperatures **(e)**, hairpin temperatures **(f)**, homo-dimer temperatures **(g)** or the GC content **(h)** were calculated either for the primer sequence including the most common nucleotides or averaged over all permutations of the final primer sequences that include degenerate nucleotides. (e-f) were calculated with primer3. Each dot represents a single primer of the respective tiled primer schemes. Dotted lines indicate the upper target cut-offs or target ranges employed by varVAMP (nt - nucleotide).

First, we analyzed the degeneracy per primer as this is highly penalized by varVAMP to keep the number of permutations minimal. Two, four and five tolerated ambiguous bases within a primer sequence can lead to a maximum degeneracy of 4, 256 and 1024, respectively. However, for the primer schemes with four and five tolerated ambiguous bases the mean number of permutations was over 10-fold lower than theoretically possible, indicating a preferential selection of primers with a low degeneracy (Fig. 3b). With the integration of ambiguous bases, varVAMP aims to minimize mismatches with the input MSA. Therefore, we analyzed for each scheme the number of mismatches against sequences in the MSA (Fig. 3c). Indeed, the large majority of build consensus sequences did not have sequence variations not covered by any primer permutations with ratHEV having higher number of mismatches likely due to the sequence variability of the MSA (Fig. 3 a). varVAMP penalizes mismatches in the last five bases of a primer’s 3’ end to ensure stable target binding. By analyzing the position-dependent mismatches of all primers in a scheme, we indeed observed that most sequences of the input MSA displayed a low frequency of mismatches at the 3’ end of the primers (Fig. 3d). Starting at the 3’ end, the number of mismatches increased in consistent waves of three nucleotides for all primer schemes except SARS-CoV-2. We hypothesized that this might be due to synonymous codon usage caused by variations in the second and third codon position^52,53^. Manual inspection of the primers’ locations in the MSAs indeed confirmed that our 3’ penalty system preferentially selected primers if their 3’ ends are located at the first and not at the second or third position of a codon. Lastly, we explored the hypothesis that the mean primer parameters of all primer permutations would lie within our target range even if they were initially calculated on the basis of the primer sequence including the most common nucleotides. We therefore calculated melting temperature, hairpin temperature, homo-dimer temperature and GC content (Fig. 3 e-h). In most cases, the mean of the primer permutations were within the target range or below the cutoff but showed a higher deviation from the optimum compared to the primer that was initially used for parameter calculation. The GC content is the least penalized parameter by varVAMP and other parameters should have a more pronounced effect on primer selection with the current settings. Indeed, the GC content was also within the target range but overall more dependent on the MSA’s GC content (Fig. 3 h).

In conclusion, the newly designed primers should recognize the majority of sequences in the initial MSA while minimizing degeneracy, overall mismatches, and 3’ mismatches.

### Full genome tiled amplicon sequencing of SARS-CoV-2, BoDV-1, HAV, PV and ratHEV

In a multi-center study with specialists for the respective pathogens, we evaluated if the newly designed primers for SARS-CoV-2, BoDV-1, HAV, PV and ratHEV were suitable for whole-genome sequencing and genome reconstruction. Similar to the HEV-3 primer schemes, we performed amplicon-based Illumina and, in the case of SARS-CoV-2 and some of the HAV samples, ONT sequencing on various samples in either singleplex or multiplex PCR reactions. Sequencing protocols and selection of samples differed due to center-specific preferences. For SARS-CoV-2, we tested the novel primer scheme in multiplex PCR reactions on a random set of respiratory patient specimens from currently circulating variants with different viral loads. Although some amplicons had a lower coverage, we were able to construct complete genomes in the majority of cases (Fig. 4a and S1). We evaluated the BoDV-1 primers in multiplex reactions on three different virus stocks that had been isolated from brains of diseased patients in 2019, 2020 and 2022^34–36^ and were cultivated on Vero cells. For all isolates we were able to recover highly covered genome sequences (Fig. 4b and S1). Only for the 2022 isolate, the last three amplicons were poorly amplified, leading to a slightly lower genome recovery (S1). For HAV, we tested the HAV-specific primers on the cell culture derived lab strain V18-35519. Illumina sequencing yielded consistent and high coverage over all amplicons independent of multi- or singleplex reactions (Fig. 4c). Next, we transferred the protocol to three different HAV-positive patient samples: genotype IB-positive feces (patient 1) and sera (patient 3) as well as genotype IA positive feces (patient 2). Finally, we sequenced another four patients sera via Oxford Nanopore: IA-positive (patient 4 and 5), IB-positive (patient 6) and IIIA-positive (patient 7). Full-genome recovery was achieved with all samples. However, for some of the amplicons we observed a lower overall coverage compared to the virus isolate (S1). Next, the PV primer scheme was tested on the Sabin 1-3 vaccine strains. Similar to our prior results, sequencing resulted in high coverage and full-genome recovery. However, we observed that the third amplicon overall under-performed in multiplex but not in singleplex reactions (Fig. 4d and S1). Lastly, we evaluated the ratHEV primers that we designed to test the limits of varVAMP given the highest sequence variability and low number of MSA sequences (Table 1). We tested either single- or multiplex PCR reactions for the two previously described isolates R63 and pt2^39,40^ (Fig. 4e). While we were able to achieve high coverage and genome recovery for the R63 isolate, we observed one and two amplicon dropouts for the pt2 single- and multiplex reactions, respectively (S1).

**Figure 4.**
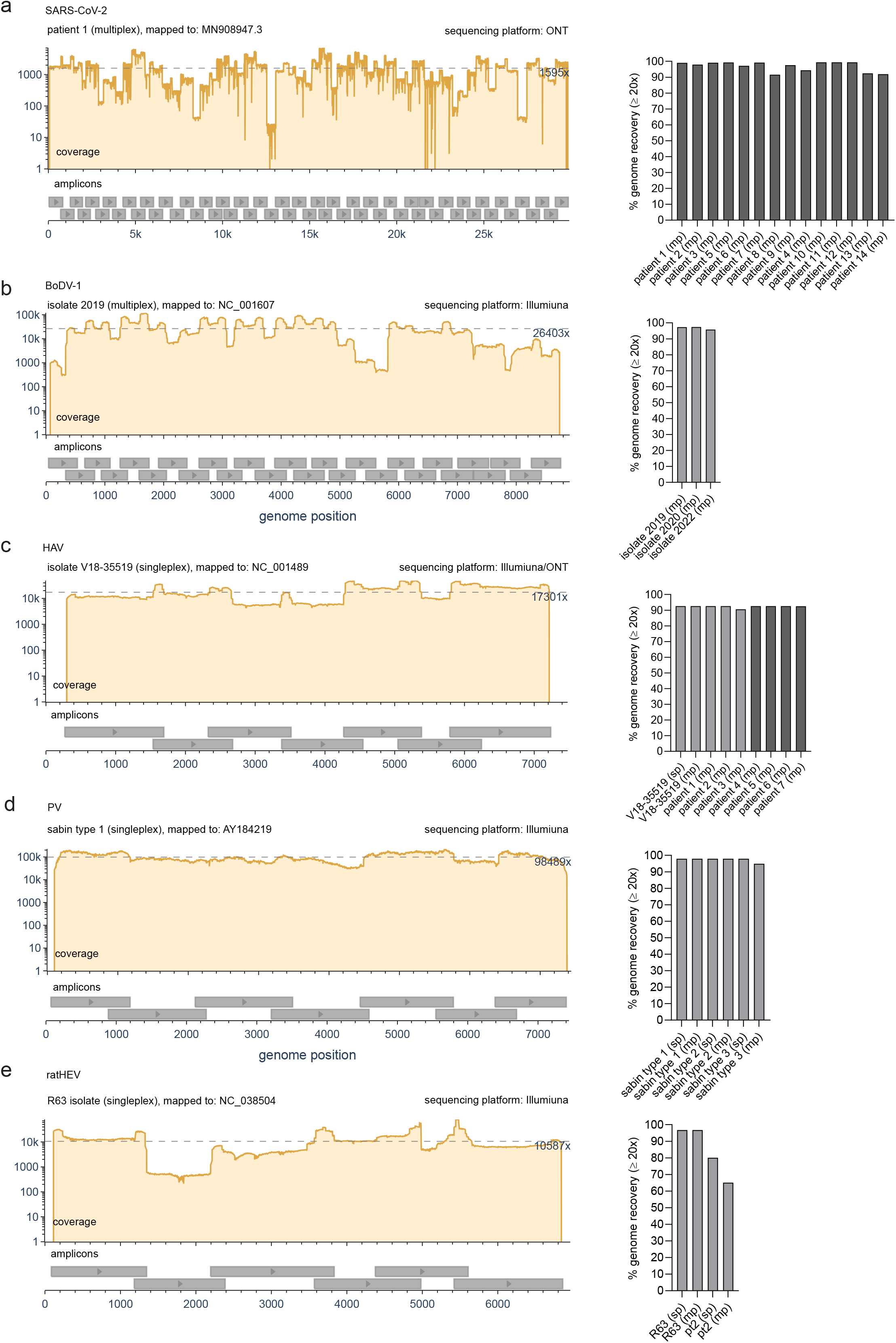
Whole genome sequencing utilizing the SARS-CoV-2, BoDV-1, HAV, PV and ratHEV primer schemes. Representative coverage plots (left) and % genome recovery (right) of the different **(a)** SARS-CoV-2, **(b)** BoDV-1, **(c)** HAV, **(d)** PV and **(e)** ratHEV samples subjected to their respective tiled amplicon whole genome sequencing workflow. Coverage plots were created with BAMdash. Dotted lines indicate mean coverages. Reference genomes used for mapping are indicated in the header of the coverage plots (individual coverage plots are given in S1). Genome recovery was calculated as % of reference nucleotides covered at least 20 fold (sp - single plex, mp - multiplex). Dark grey bars - ONT generated data, light grey bars - Illumina generated data.

We systematically evaluated the coverage and amplicon recovery for all primer schemes and samples (Fig. 5a). Most amplicons performed in a sample-dependent manner but in some cases multiplex performance was intrinsic to specific amplicons as exemplified by the third amplicon of the PV scheme (Fig. 4c). As all multiplex reactions across tested schemes were performed with equimolar primer concentrations, we hypothesized that the performance could be improved by balancing primer concentrations. Therefore, we adjusted the molarity of the PV primers in two consecutive rounds as a proof-of-concept for further wet-lab optimization. For the final iteration, we achieved a coverage for all Sabin strains that was comparable to the respective singleplex reactions (S2). Analogous to the mismatch analysis with the input MSA (Fig. 3c), we also examined how many nucleotide mismatches between the primers and their target regions are present in our sequencing results (Fig. 5b). The primer target regions of the new sequences showed up to two mismatches to the degenerated primer sequences with the majority having no mismatches. Similar to the amplicon performance, the number of mismatches were mostly sample-dependent. As the samples used for evaluation were selected in the respective centers based on availability and not sequence diversity, we tested if this could have produced an unintended selection bias towards specific viral strains. Therefore, we evaluated if the novel consensus sequences of each scheme have a variability that is comparable to that of the respective input alignments. Pairwise sequence identities of these small datasets were highly similar or lower to that of the alignment with only the newly produced sequences for HEV cluster 4 showing a significantly higher mean pairwise sequence identity of 7 %, respectively (Fig. 5c), indicating a slight bias for higher conserved sequences.

**Figure 5.**
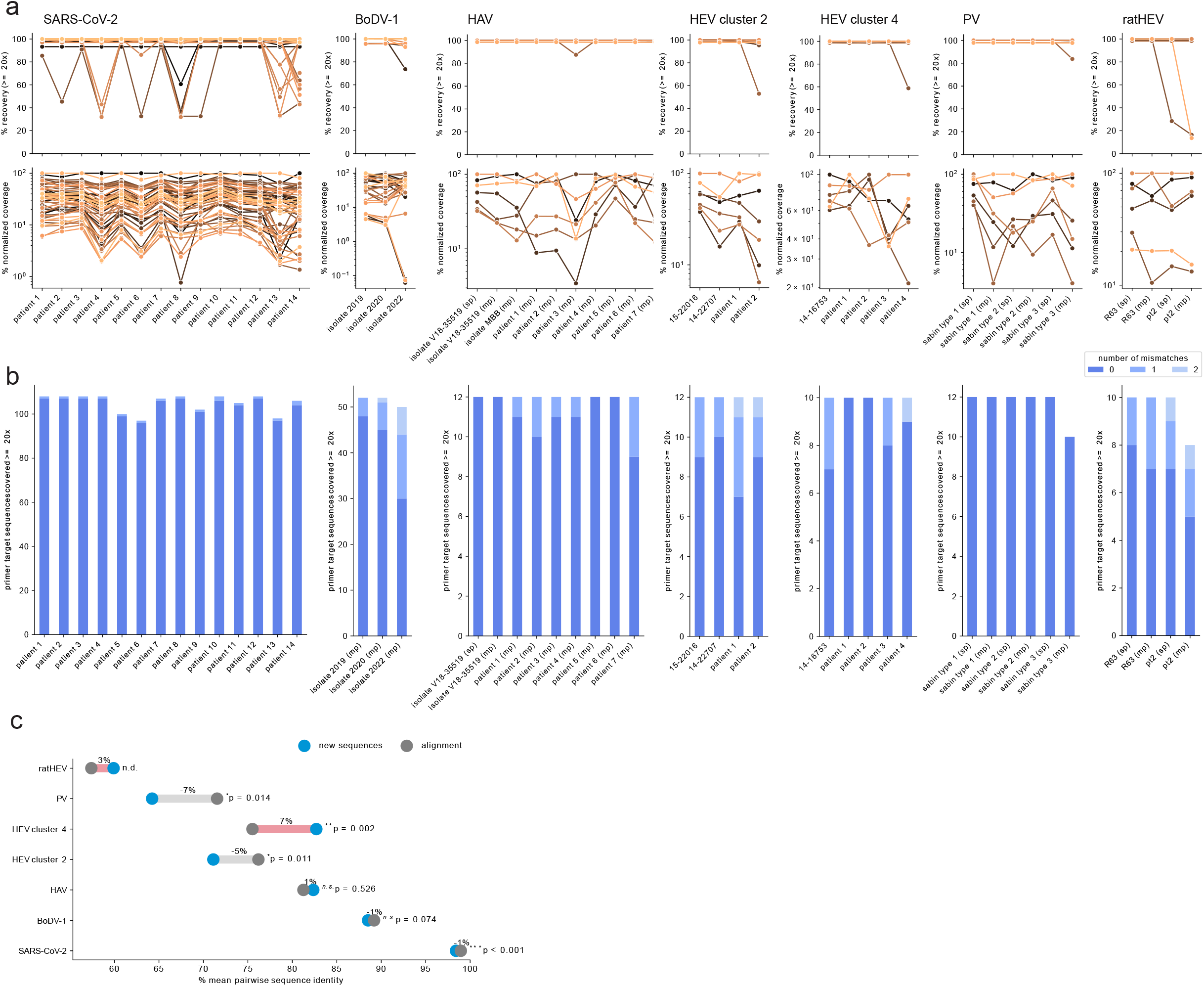
Amplicon performance and mismatch analysis. **(a)** For each sequencing result using the virus specific primer scheme the amplicon recovery (upper panel) and normalized coverage (lower panel) was calculated. Each color represents an individual amplicon tracked over different samples. Amplicon recovery was calculated as % of reference nucleotides covered at least 20 fold between the genomic start and stop position of the individual amplicons. For the normalized amplicon coverage, the mean coverage was calculated for each amplicon and normalized to the highest covered amplicon of each scheme (set to 100). **(b)** For each sequencing result, each primer binding region was analyzed for the number of mismatches not covered by any permutation of the corresponding primer sequence. Mutations were only considered if their variant frequency was >= 0.7. Primers were excluded from the analysis if any primer binding position was not covered at least 20-fold. **(c)** Dumbbell plot showing the pairwise identities of the newly generated fasta consensus sequences (blue dot) or the sequences of the varVAMP input MSA (dark gray dot) of each respective primer scheme. Light gray and red lines indicate the percent pairwise identity increase or decrease, respectively. Significance was calculated with a Welch’s *t*-test (n.d. - not determined as n < 3, n.s. - not significant, *: p ≤ 0.05, **: p ≤ 0.05).

In summary, all primer schemes were suitable for tiled amplicon Illumina or ONT sequencing and resulted in highly covered full-genome sequences by applying center-specific sequencing protocols and bioinformatic pipelines initially developed for tiled amplicon sequencing of SARS-CoV-2.

### Design and wet-lab evaluation of PV qPCR primers designed with varVAMP

The WHO gold standard for PV detection is based on time- and resource-consuming virus cultivation. Molecular detection by qPCR is available but was designed for virus isolates propagated in cell culture^54^. Moreover, primers and probes display a high level of degeneracy, decreasing the assay’s sensitivity and increasing the risk of unspecific non-viral amplification products for other specimen sources like wastewater samples. Therefore, we used varVAMP to design PV serotype-specific assays. Optimal primer annealing temperature of each assay was tested with a gradient PCR ranging from 56 - 64 °C. All annealing temperatures resulted in the expected product size with 59 °C showing the lowest abundance of unspecific products. PV serotype specific RT-qPCR assays were performed in a serial dilution experiment between 10^−1^ and 10^−8^ with RNA extracted from viral supernatant. For all three types, PV detection was achieved up to a dilution of 10^−7^ (Fig. 6 a-c). Absolute quantification analysis based on the dilution series calculated an efficiency value (PV1 = 1.88, PV2 = 1.87, PV 3 = 1.90) close to 2 corresponding to a perfect amplification reaction. Cross-specificity testing showed no detection between the Sabin strains indicating that the designed primers and probes are highly specific for their respective PV serotype, despite primer and probe degeneration.

**Figure 6.**
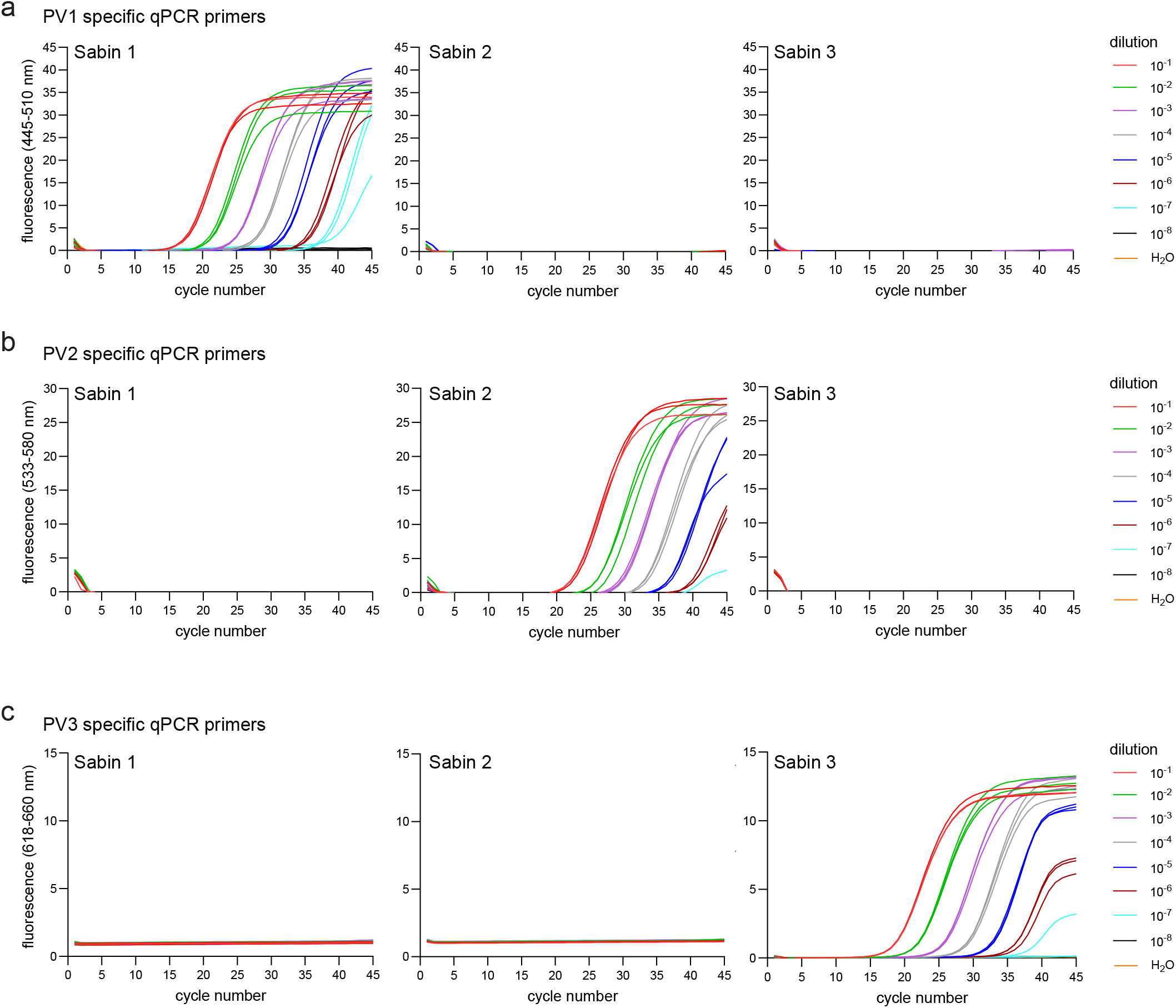
Specificity and sensitivity of the novel PV qPCR schemes. qPCR primers specific for **(a)** PV1, **(b)** PV2 and **(c)** PV3 were tested on serial RNA dilutions extracted from viral supernatants of Sabin 1, Sabin 2 and Sabin 3 infected cell cultures (n=3). The fluorescence was measured during the extension step with three different channel setups respective to the probe fluorophore (FAM = 465-510 nm, JOE = 533-580 nm, CY5 = 618-660 nm). Amplification curves were analyzed using the Roche LightCycler 480 II device software.

## DISCUSSION

Here, we describe varVAMP, a command-line software tailored to pan-specific primer design for highly variable MSAs. Importantly, varVAMP is available through various bioinformatic repositories and has also been deployed to Galaxy Europe (usegalaxy.eu), a web-based platform for bioinformatic data analysis^55^. On the basis of the input MSA, varVAMP generates consensus sequences and analyzes them for the presence of potential primer sequences. From the subsequent pool of found primers, varVAMP chooses optimal amplicons for specific molecular techniques such as tiled sequencing or qPCR and introduces degenerated nucleotides into primer sequences to compensate for sequence variations.

To demonstrate varVAMP’s applicability, we used the software to design and evaluate primers for tiled sequencing schemes of SARS-CoV-2, BoDV-1, HAV, HEV, PV and ratHEV and for qPCR of PV. With the exception of SARS-CoV-2, these pathogens have a high genomic diversity restricting conventional primer design. While varVAMP provides a fully automated solution, selecting appropriate reference data prior to primer design can pose a challenge. The core concept of varVAMP is based on the assumption that the sequences in the input MSA reflect the majority of variations within the viral genome. However, in sequence data repositories, particular in the case of viral sequences, there can be a lack of associated metadata, the presence of recombinant sequences, bias towards sequencing labs, geographic biases towards circulating strains or an underrepresented amount of recently discovered or understudied viral pathogens^56,57^. Therefore, a prior careful data selection is a key requirement for a successful primer design and might require the additional use of clustering algorithms and phylogenetic assessment tools^29,30,58,59^. This was most prominently observed for our ratHEV tiled sequencing scheme. Here, we could only successfully generate full length sequences for the R63 but not for the pt2 isolate. The ratHEV primer design was challenging as ratHEV full-length sequences are highly variable and scarce in NCBI’s genbank. In such cases, PCR reactions might not be successful for viral genomes that are only distantly related to the majority of sequences in the input MSA. The pathogenicity of ratHEV for humans has been only recently identified^60^. We expect an increase in reported sequence data over the coming years, which will help to overcome current limitations. The second challenge for varVAMP users is the proper selection of input parameters. While varVAMP offers an automated solution, it often requires multiple rounds of manual optimizations that should, however, be computationally inexpensive.

It is worth mentioning that we did not compare our primer design suite to other primer design software, as we are not aware of other tools that handle similar sequence variability while being able to design degenerate primers for tiled sequencing or qPCR. Primalscheme was highly successful during the SARS-CoV-2 pandemic and has been also used to design tiled amplicon schemes for viruses such as West-Nile or Monkeypox virus^61–63^ but limits the variability of the input alignment at 5%^10^. Despite the fact that varVAMP adapted some of primalscheme’s primer assessment functionality, a head-to-head comparison is not possible as varVAMP was not designed as an alternative but as a solution for highly variable MSAs.

We did observe sample-dependent and independent amplicon performance. The molecular reasons for sample-dependent PCR performance can be diverse. We show that primers lead to amplification even if there are one or two mismatches between primer and target sequence. However, if more mismatches are present, poorer amplification or complete dropouts could be the result. For patient samples, we used archived material with varying storage times and viral loads that both could have impacted PCR amplification. Another problem can be the presence of sample-specific non-viral nucleic acid that could provide additional primer binding sites and result in unspecific amplification. To address this issue, varVAMP provides a BLAST module that allows users to check for potential off-target effects with a custom database. Sample-independent amplicon performance is likely a PCR optimization issue. Compared to popular SARS-CoV-2 schemes^64^, we have not optimized the PCR conditions, nor the primer pooling ratios. That was reflected by varying amplicon performance in multiplex reactions. To give an example of how to further optimize the multiplex PCRs, we adjusted the primer concentrations of the PV scheme and show this can lead to a more balanced overall coverage.

All primer schemes described here have been developed because of a methodology gap. HEV phylogeny and sub-typing is mostly restricted to Sanger sequencing of ORF2 fragments thereby neglecting viral evolution in the remaining genome^65,66^. Current methodology to generate highly covered HEV full-genome sequences via Illumina or Nanopore sequencing requires costly RNA-Seq protocols using hybridization probes^67,68^ or powerful sequencing machines. Our newly developed primer schemes could simplify sequencing and aid, for example, analyses of Ribavirin resistance-associated mutations that can develop in immunocompromised HEV patients^69^. For PV, molecular assays used for detection and intratypic differentiation of serotypes are well suited for cell-culture isolates^70^. However, these techniques are laborious and time-consuming. qPCR and amplicon-based NGS protocols for the rapid analysis of PV from patient and environmental samples are imperative for fast public health decisions. There are protocols for high-throughput sequencing of PV but similar to HEV they are restricted to a conserved part of the enterovirus genome^71^. Our novel methods for PV detection by qPCR and whole-genome sequencing could not only benefit existing surveillance programs but might also lay the foundation for wastewater surveillance strategies within the global PV eradication program^72,73^.

In conclusion, the varVAMP pipeline was developed because primer design on the basis of highly variable MSAs is difficult, time consuming and there are no automated solutions for qPCR and tiled sequencing. The designed and validated primer schemes for the different viruses are not only a proof-of-concept for varVAMP’s applicability but have been developed because they could directly benefit viral diagnostics and epidemiology. Laboratories that have already established SARS-CoV-2 sequencing pipelines or in-house qPCR protocols should be able to adapt their methodologies to these new primer schemes with only minor modifications.

## DATA AVAILABILITY

varVAMP v.1.2.0 and BAMdash v.0.2.4 are open source and available at https://github.com/jonas-fuchs/varVAMP (DOI: 10.5281/zenodo.11125498) and https://github.com/jonas-fuchs/BAMdash (DOI: 10.5281/zenodo.10804160). The Galaxy version of varVAMP is available at https://usegalaxy.eu/root?tool_id=toolshed.g2.bx.psu.edu/repos/iuc/varvamp/varvamp/. The code and data to reproduce the figures is available at https://github.com/jonas-fuchs/varVAMP_in_silico_analysis (DOI: 10.5281/zenodo.10942525). All input multiple sequence alignments and varVAMP outputs for primers that have been evaluated in this study are available at: https://github.com/jonas-fuchs/ViralPrimerSchemes (DOI: 10.5281/zenodo.10562882). Raw sequencing data has been deposited at ENA under the study accession number: PRJEB74744.

## AUTHOR CONTRIBUTIONS

JF, MH, MS, WM, MP conceptualized the project. JF wrote the manuscript. JF developed varVAMP with help from WM. WM and BG deployed varVAMP to CONDA, DOCKER and Galaxy. TK critically evaluated the algorithms employed by varVAMP. MS and JF selected the input data for primer design. JF and JKl designed primers with varVAMP. JKl, MS, JKr, CW, LE, LJ, AM, CB, MB, JP, RJ, JW, JS, CM, SB, SS generated and evaluated the experimental data. JF, NB, JKl and MH performed the bioinformatic analyses. JF performed phylogenetic analyses and the *in silico* primer evaluation. All authors reviewed and edited the manuscript.

## ACKNOWLEDGEMENTS

We would like to thank Dr. Freya Fleckenstein for many suggestions for the code base of varVAMP and for help with the mathematical part of the methods section. We would also like to thank Dr. Zsolt Ruscics for his valuable and critical input at the early stages of varVAMP development. We thank Ashkan Ghassemi for his help in preparing representative SARS-CoV-2 genomes and Dr. Christian Blumenscheit for help in ordering the SARS-CoV-2 varVAMP primers. We also appreciate the effort of the submitting laboratories for the procurement and provision of SARS-CoV-2 genomes via the German Electronic Sequence Data Hub and those laboratories providing SARS-CoV-2 positive samples via the IMS-SC2 network. We acknowledge the Sequencing Core Facilities of the Genome Competence Centre at the Robert Koch Institute for providing excellent sequencing services for the PV and SARS-CoV-2 samples, and Aaron Houterman for his excellent technical assistance. We would also like to thank Prof. Dr. Otto Haller for his helpful comments on the manuscript.

## TABLE AND FIGURE LEGENDS

**S1. Coverage plots for all sequencing results**. Coverage plots of the different SARS-CoV-2, BoDV-1, HAV, PV and ratHEV samples subjected to their respective tiled amplicon whole genome Illumina sequencing workflow. Dotted lines indicate mean coverages. NCBI accession numbers of the reference sequences used for mappings are indicated in the headers. Coverage plots were created with BAMdash.

**S2. Primer balancing for PV whole genome sequencing**. The PV tiled primers initially used in equimolar concentrations for multiplex reactions were balanced in two consecutive rounds based on prior results and then the balanced primers were used in multiplex PCR reactions for Sabin 1-3 and the amplicons subjected to Illumina sequencing. The respective concentrations for each iteration are given above the coverage plots (blue arrow - increase in molarity, gray arrow - no change in molarity, red arrow - decrease in molarity). Dotted lines indicate mean coverages. NCBI accession numbers of the reference sequences used for mappings are indicated in the headers. Coverage plots were created with BAMdash.

