## Supplementary figures and images for "varVAMP: automated pan-specific primer design for tiled full genome sequencing and qPCR of highly diverse viral pathogens"

### Supplementary Figures

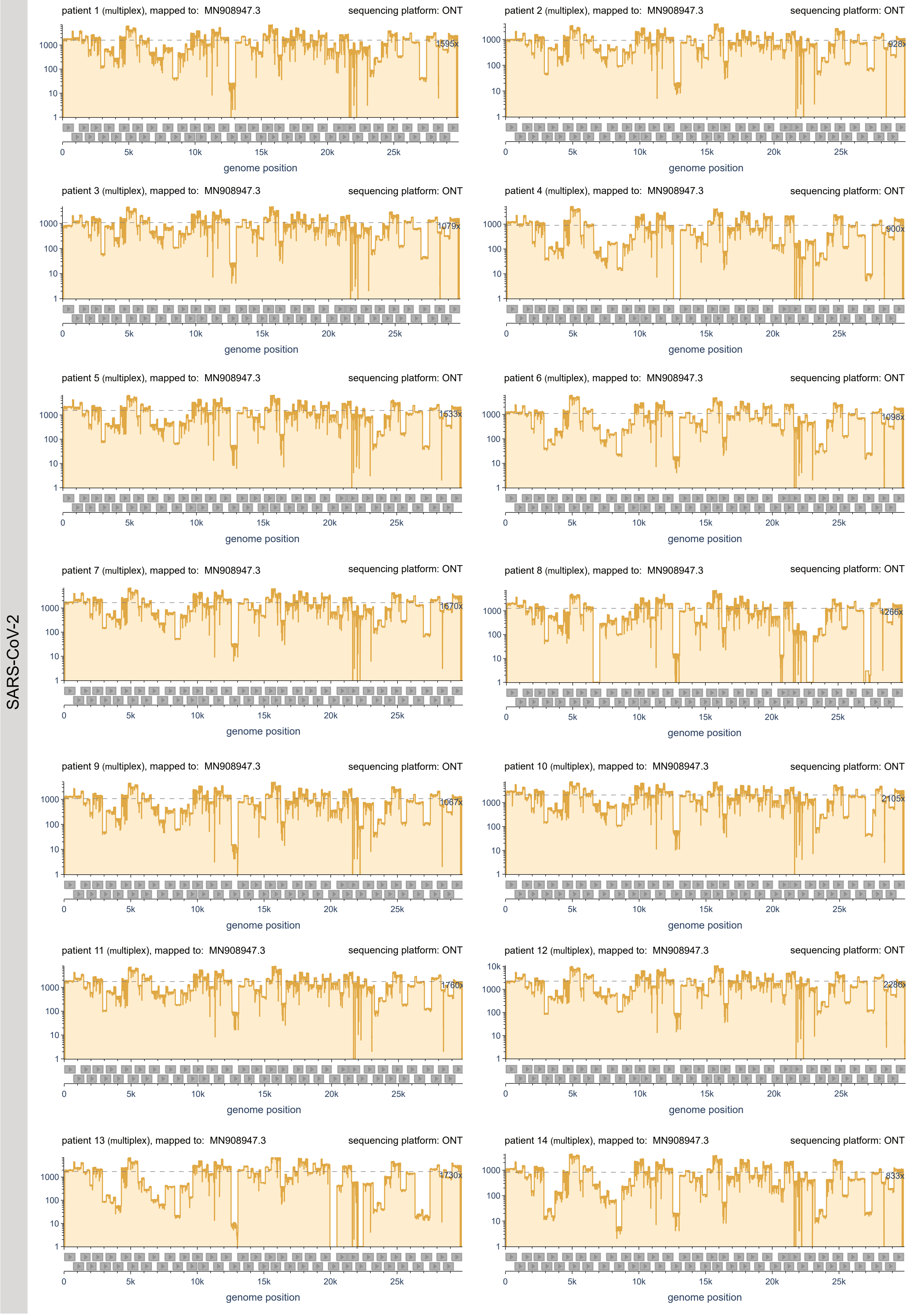

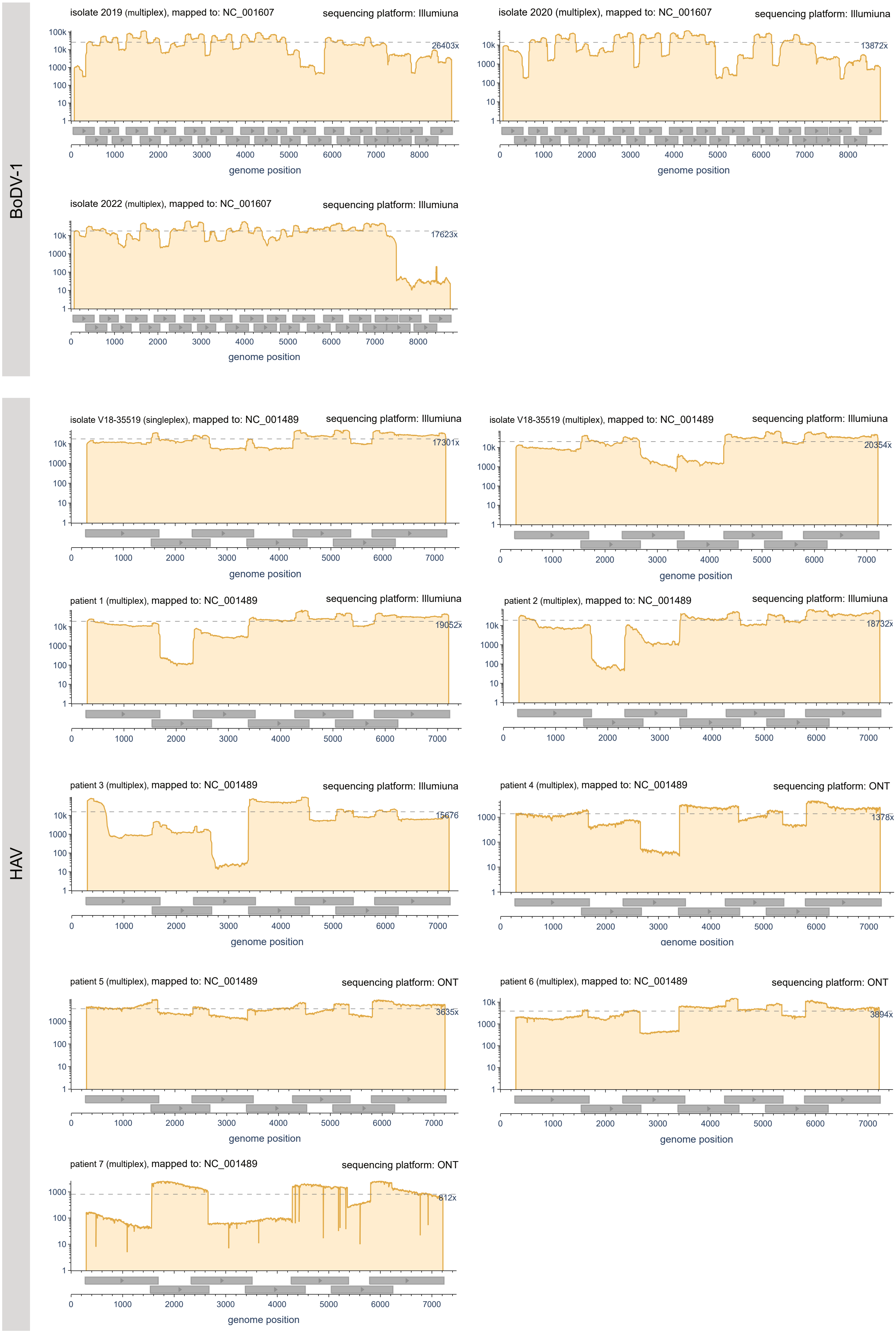

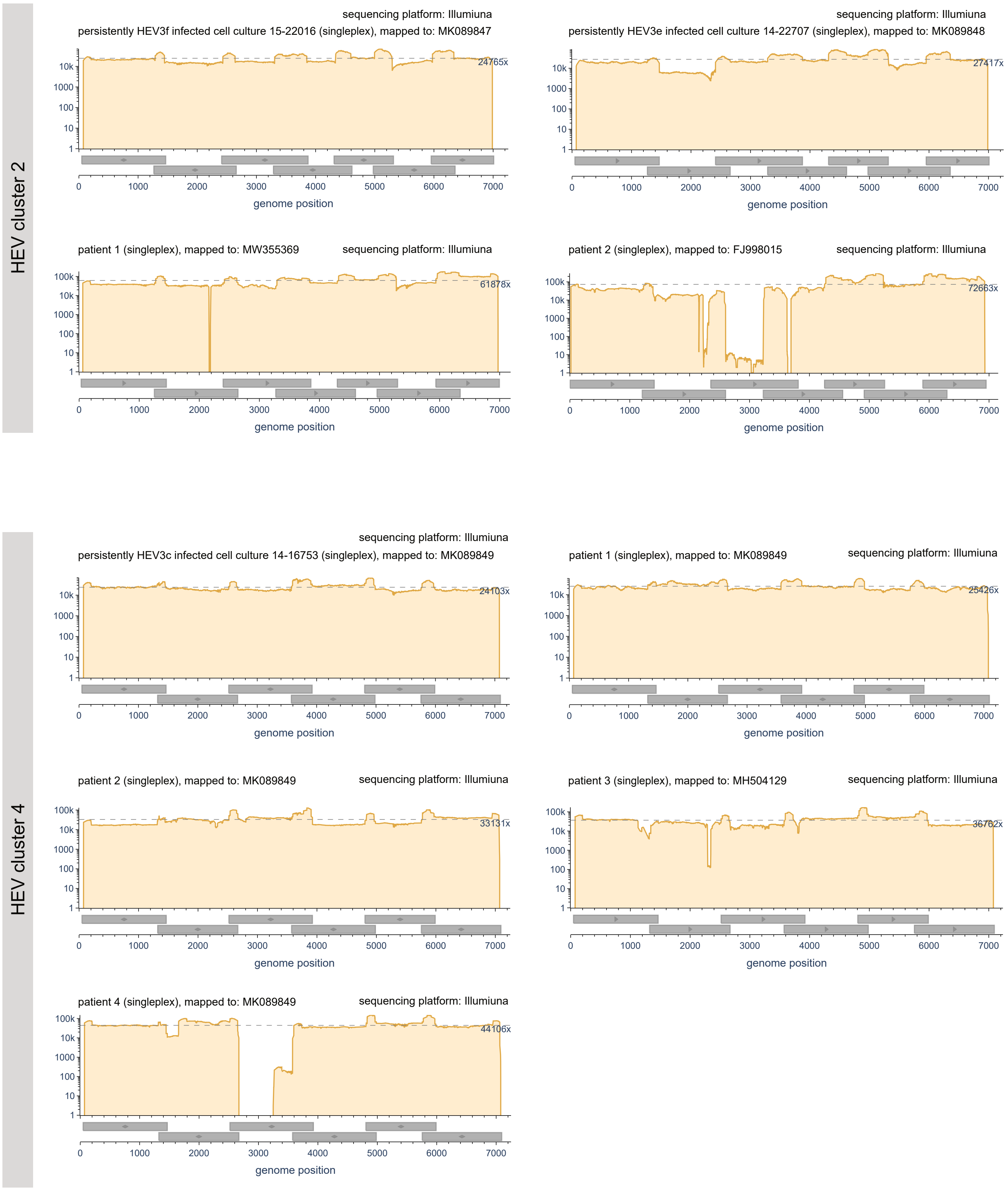

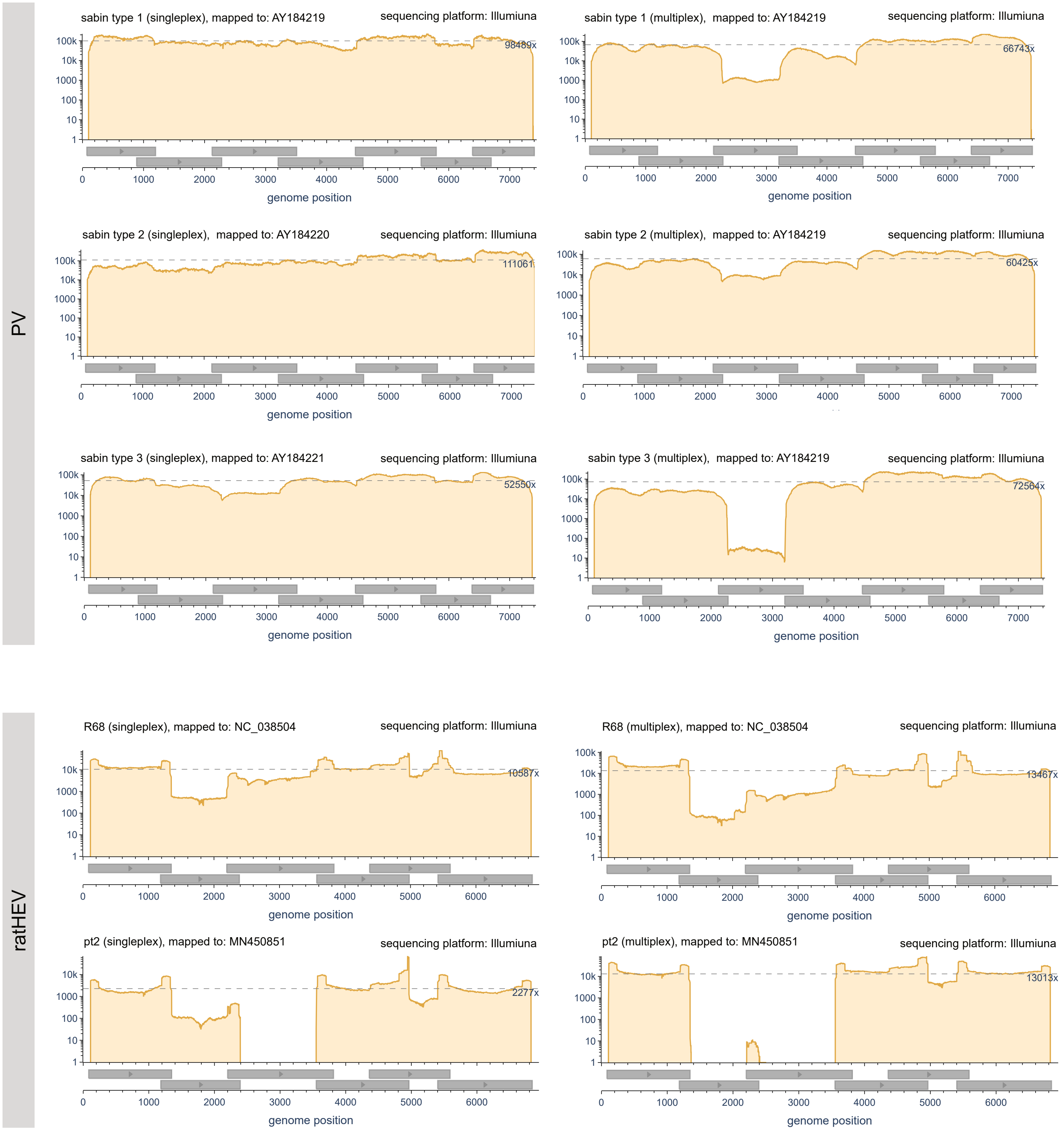

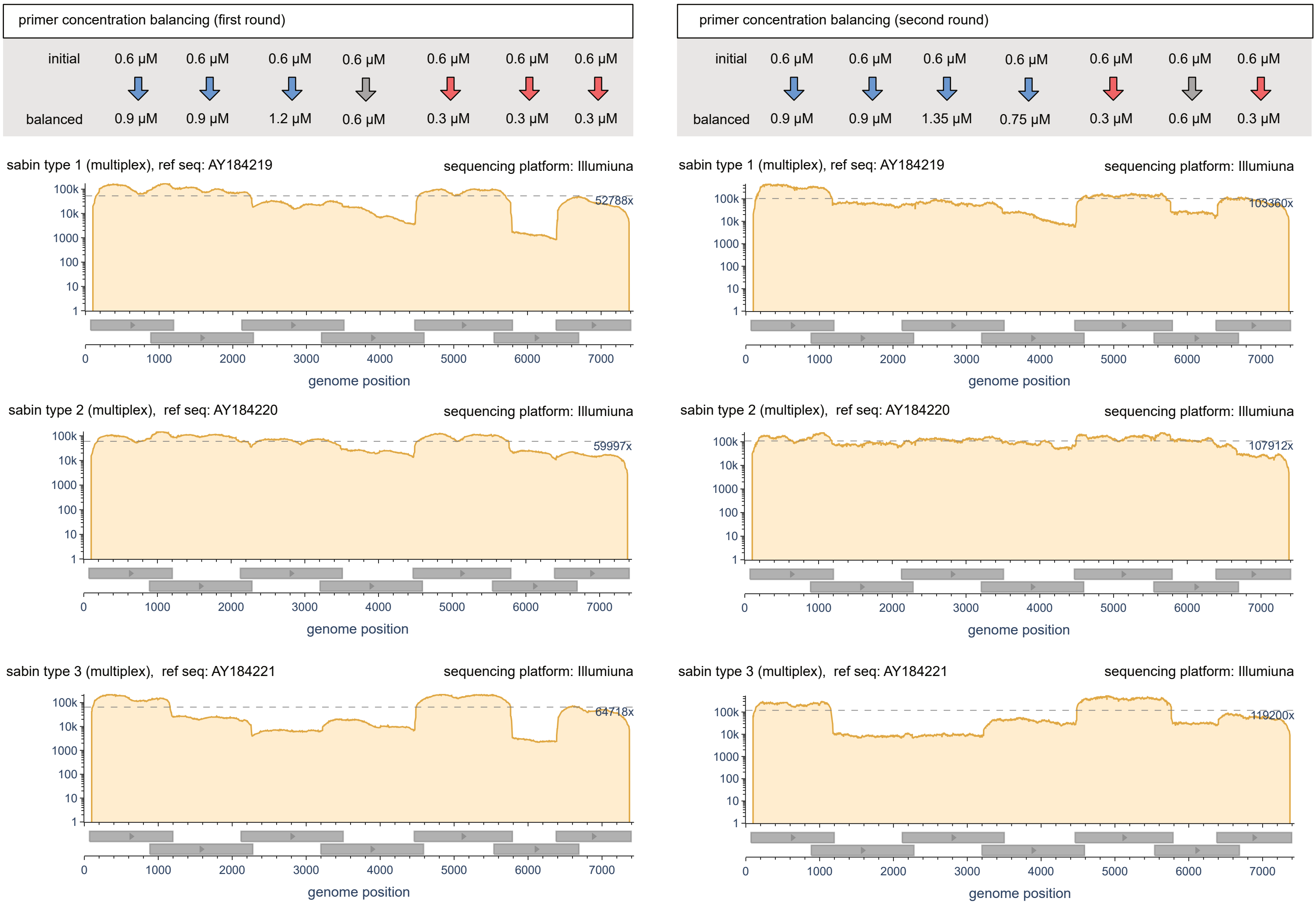
